# Dynamic uterine microenvironment drives endometrial adenocarcinoma carcinogenesis and progression

**DOI:** 10.1101/2025.08.11.669782

**Authors:** Lesley B. Conrad, Shiwei Yin, Bingru Feng, LaShelle King, Brenna Hobson, Katelyn Andersen, John Coté, Robin Farias-Eisner, Jun Xia, Yusi Fu

## Abstract

Endometrial cancer (EC) development is driven by the interaction between the tumor and the microenvironment. However, the immune microenvironment dynamics during this process are not clear. Here, we applied single-cell RNA sequencing (scRNA-seq) to uterine blood samples collected at hysterectomy from 15 individuals encompassing four groups—benign controls, endometrial intraepithelial neoplasia (EIN), the transition from EIN to carcinoma, and EC. Uterine blood, obtained without prior enrichment, provides a liquid biopsy of the local tumor milieu, enabling high-resolution profiling of both immune and stromal cells in a minimally invasive manner. Our analysis revealed simultaneous immune and stromal remodeling in early premalignant lesions. Notably, even in the EIN, we observed significant immune cell reprogramming alongside the emergence of protumorigenic stromal-epithelial interactions. Importantly, we discovered predictive single-cell transcriptomic signatures derived from neutrophils that stratify patients according to disease state, highlighting the potential of tumor-educated innate immune cells as biomarkers. These findings yield candidate cellular and molecular signatures, particularly from neutrophils, that could enhance early EC detection and guide therapeutic strategies. Our work provides a proof-of-concept for leveraging local liquid biopsies in single-cell oncology, offering new insights into EC initiation and paving the way for noninvasive diagnostics on the basis of single-cell transcriptomic profiles.

## Introduction

Endometrial cancer (EC) is the most prevalent gynecologic malignancy in developed countries, with approximately 65,000–70,000 new diagnoses and over 12,000 related deaths each year in the United States^1^. Notably, EC is one of the few cancers with increasing trends in both incidence and mortality^2^. The primary precursor lesion to endometrioid EC, the most common EC histologic subtype^3^, is endometrial intraepithelial neoplasia (EIN), a premalignant condition characterized by dysplastic, clonal proliferation of endometrial glands that confers a growth advantage over the surrounding stroma^4^. These atypical hyperplasia lesions often progress to or coexist with or progress to EC if left untreated, underscoring their clinical significance in endometrial carcinogenesis^5^. Despite substantial efforts to map the genomic and transcriptomic alterations in endometrial carcinoma^6,7^, our understanding of the coevolution between the neoplastic endometrial epithelium and the local immune microenvironment remains limited. There is a knowledge gap in how immune cells and epithelial cells within the uterus interact with each other and influence each other during the progression from the normal endometrium to EIN and ultimately to invasive carcinoma.

Emerging evidence points to the important role of the immune system in tumor initiation and progression^8^, motivating approaches that jointly examine tumor and immune compartments. In this study, we introduce a unique methodology that captures both endometrial epithelial and immune cells from the uterine environment by analyzing uterine blood samples rather than relying solely on traditional tissue biopsies. Uterine blood collected from the uterine cavity contains cells from the local tumor microenvironment, offering a refined and sensitive sampling method to detect concurrent changes in both the endometrial lining and immune cell populations during the transition from a healthy endometrium through the EIN to the EC. By directly comparing analyses of uterine blood as a liquid biopsy to those of endometrial tissue biopsies^7^, we demonstrate that uterine blood provides a more localized snapshot of the uterine microenvironment. This approach increases the sensitivity for capturing cellular and molecular changes, and it yields a broader array of potential biomarkers across multiple cell types that are predictive of EC development on the basis of their expression signatures.

By untangling the complex cellular heterogeneity of a tumor and its microenvironment, we aim to develop an integrated molecular signature that incorporates both tumor-intrinsic genomic alterations and tumor microenvironment (TME) immune features for the early detection of malignancy. In our analysis, we profiled the single-cell transcriptomes of 15 individuals across the spectrum of endometrial carcinogenesis, capturing 35 distinct cell types and a total of 46,583 single cells spanning four key time points from the normal endometrium to the EIN to the EC. This rich dataset enables us to pinpoint stage-specific gene expression changes and cellular interactions that characterize the stepwise progression of EC.

## Results

### A single-cell atlas reveals immune and stromal remodeling during EC development

To simultaneously delineate the changes in the endometrium and uterine immune profiles during EC development, we collected uterine blood during hysterectomy from 15 individuals spanning the evolutionary trajectory of endometrial cancer and categorized them into 4 groups: cancer-free, preinvasive (EIN), invasive transition, and EC (Fig. 1a), with demographic and clinical profiles provided in Supplemental Table 1. Cancer-free samples are those with a benign endometrium who underwent hysterectomy for benign indications; EIN samples represent endometrial cells that start gaining overgrowth ability as a precursor lesion to invasive carcinoma; invasive transition samples are EIN on biopsy but cancer on hysterectomy pathology, representing samples with endometrial cells just gaining invasiveness, and finally EC patients. All EC samples included were uterine-confined carcinomas (International Federation of Gynecology and Obstetrics (FIGO)^9^ stage I), except one sample with extra-uterine disease (FIGO stage III). This sample was included in our analysis due to the similarity of the uterine immune profiles to uterine-confined samples. These four groups represent the transition from normal endometrium to development of invasive carcinoma and aid in identifying important biomarkers and molecular changes during EC development.

**Figure 1.**
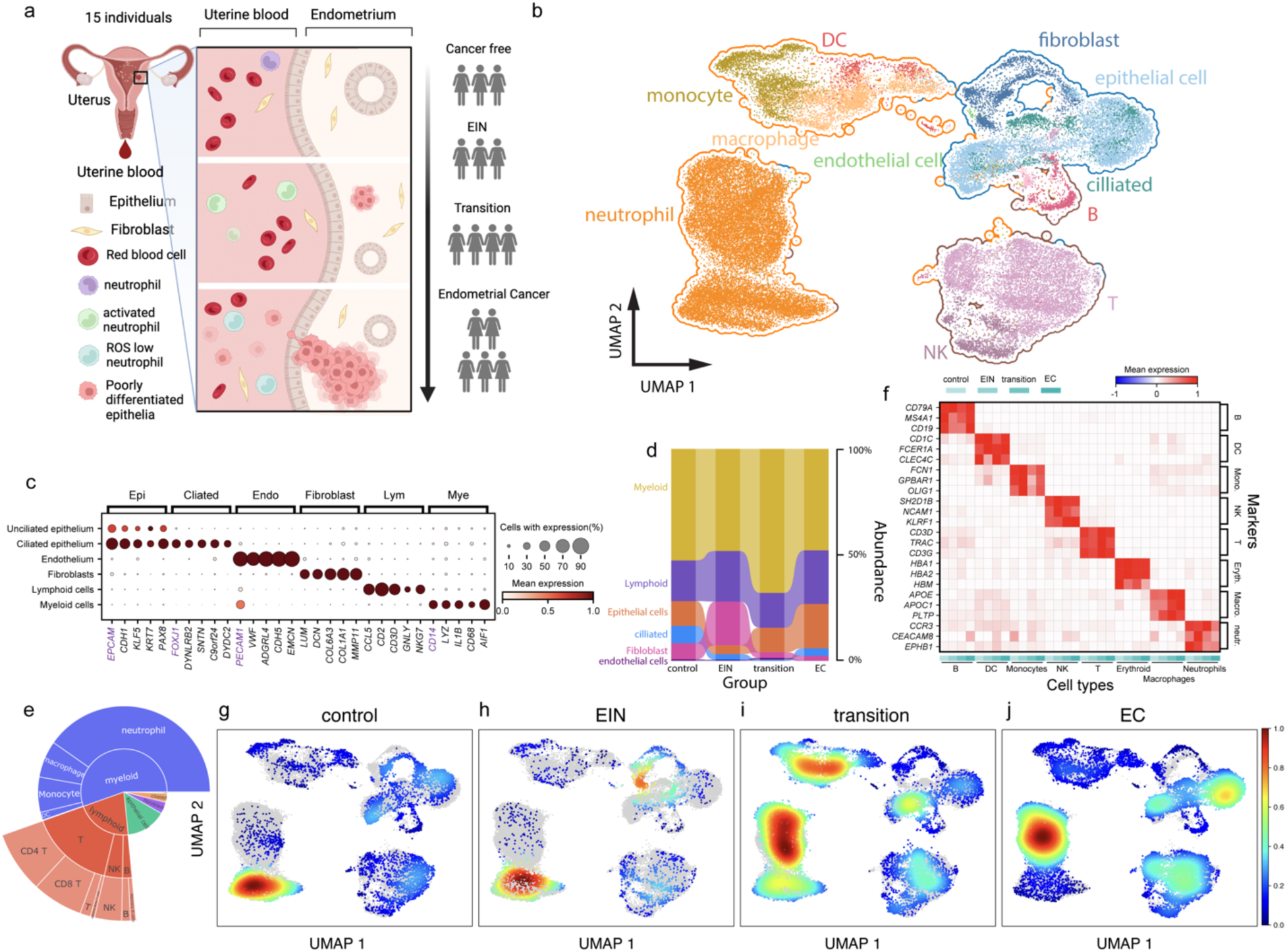
A single-cell atlas of the endometrium and uterine immune environment during EC development. (a) Schematic overview of the study design. Uterine blood was collected at the time of hysterectomy from 15 individuals spanning four stages of EC evolution: cancer-free, preinvasive endometrial intraepithelial neoplasia (EIN), invasive transition, and EC. Demographic details are provided in Supplementary Table 1. (b) Uniform manifold approximation and projection (UMAP) of single-cell RNA-seq profiles from uterine blood, illustrating major cell types (myeloid, lymphoid, epithelial, fibroblast, smooth muscle, and endothelial). (c) Heatmap of known marker gene expression across the identified cell clusters, confirming their annotations. (d) Relative abundance of the main cell types in each disease stage, showing a decrease in fibroblasts and an increase in epithelial cells with progression toward EC. (e) Subclustering of immune cells (myeloid: neutrophils, monocytes, macrophages, and dendritic cells; lymphoid: CD8^+^ T, CD4^+^ T, NK, and B cells). (f) Dot plot of canonical markers distinguishing each immune subpopulation. (g–j) Changes in immune-cell composition across the four stages, revealing distinct shifts in myeloid and lymphoid subsets during EC development.

We performed single-cell RNA-seq on uterine blood cells from each participant to identify the molecular changes in different cell types during this cancer evolution process in both the immune system and the endometrium at the single-cell level. From the uterine blood, we were able to identify the major cell types that exist in the endometrium and endometrial cancer tissue^7,10^, including fibroblasts, the epithelium (ciliated and unciliated), smooth muscle cells, and endothelial cells, and the vast majority of the cells were immune cells, including myeloid cells and lymphoid cells, in all samples, with similar gene numbers and unique molecular identifier (UMI) counts for each cell type regardless of the group of individuals (Fig. 1b, Supplemental Fig. 1a). The expression pattern of the cells matched the known marker and previously published tissue biopsy results^7,10^ (Fig. 1c), validating the annotation of our dataset, and additional marker genes from our data revealed more tissue-specific expression patterns (Supplemental Fig. 1b). We identified similar changes in the abundances of the major cell types present in the endometrium, as previously reported from solid tissue^7^: fibroblasts decline significantly, whereas epithelial cells expand progressively as the disease progresses from EIN to invasive carcinoma (Fig. 1d, Supplemental Fig. 1c). However, the overall composition of our liquid-biopsy samples differs from that of tissue samples. Uterine-blood collections are inherently enriched for immune cells and therefore contain proportionally fewer epithelial and stromal cells (Supplementary Fig. 1d).

To identify more detailed changes in immune cells, with machine learning-based algorithms (cello^11^, CellTypist^12^), we further identified subclusters of immune cells and identified neutrophils, monocytes, macrophages and DCs among myeloid cells and CD8+ T, CD4+ T, NK and B+ cells among lymphoid cells (Fig. 1b, e), with distinct expression patterns matching known markers for each cell type regardless the group of individuals belonging to the cells derived from (Fig. 1f). Further fine annotation revealed a total of 35 distinct cell types. With this fine map of the single-cell atlas during EC development, we identified distinct transcriptomic and subpopulation changes between different groups of our samples (Fig. 1g-j) for different types of immune cells, which has not been observed previously because of the lower ratio of immune cells in the cancer tissue^7^. Our data revealed significant cellular and molecular changes during cancer development in both the endometrium and the uterine immune environment.

### Uterine blood reveals epithelial origins and genomic instability during EC progression

We first sought to validate that uterine blood sampling accurately captures the same endometrial cell populations and molecular alterations observed in conventional tissue biopsies. From our single-cell dataset, we identified both glandular and luminal unciliated epithelial cells, each displaying expression profiles consistent with known lineage markers^10^ (Supplemental Fig. 2a). Within this epithelial compartment, we discovered a distinct subcluster characterized by high expression of oncogenic markers previously reported in tissue-based studies^7^ (Supplemental Fig. 2b). Using RNA velocity-based pseudotime analysis^13^, we observed a directional trajectory from a subset of unciliated glandular cells to this oncogenic epithelial subpopulation (Fig. 2a), suggesting that unciliated glandular cells likely serve as the origin of these putative malignant cells. These findings align with evidence from EC tissues, which similarly implicates the unciliated glandular epithelium as a key source of malignant transformation^7^. Collectively, these data demonstrate that uterine blood sampling is a feasible and informative alternative to tissue biopsy for detecting cellular and molecular changes in the endometrium.

**Figure 2.**
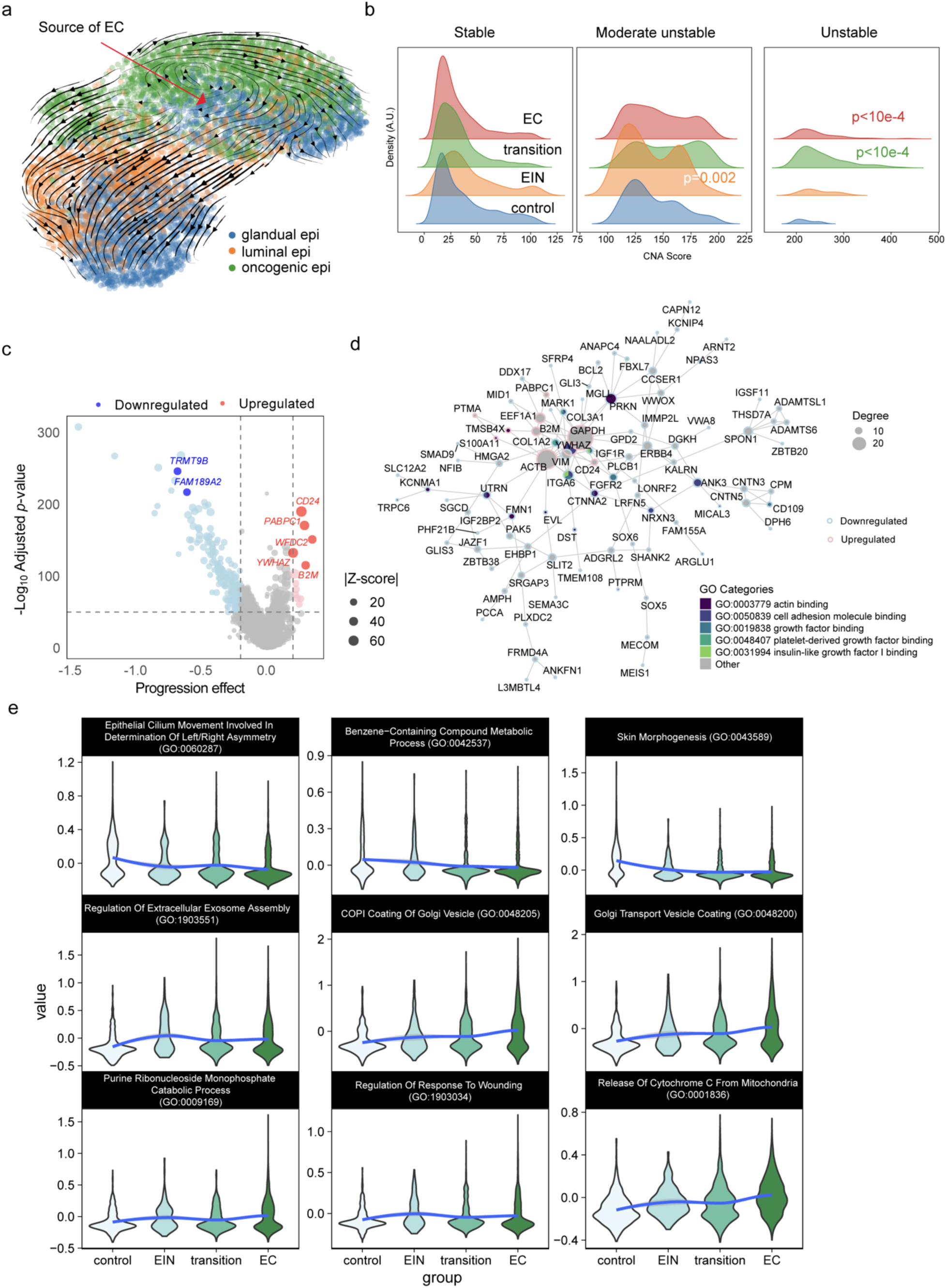
Uterine blood captures epithelial transformation and genomic instability, which are reflective of EC development. (a) RNA velocity analysis revealed a directional trajectory from glandular epithelial cells (blue) toward an oncogenic epithelial subpopulation (green), suggesting that glandular cells are likely the origin of malignant transformation in EC. (b) CNA burden-based classification of epithelial cells into genome stable (blue), moderately unstable (orange), and unstable (red) categories. The proportion of unstable epithelial cells increases progressively from control to EC, reflecting increased genomic instability during tumor development. (c) Volcano plot showing significantly upregulated (red) and downregulated (blue) genes across EC progression, based on the regression coefficient and adjusted p value. Key cancer-related genes are labeled. (d) Gene network analysis of significantly dysregulated genes identified hub genes (large nodes) with high connectivity. Node color indicates upregulation (red) or downregulation (blue); size indicates the Z score; edge thickness represents the degree of interaction. The functional enrichment of GO molecular function terms is shown. (e) Violin plots showing the expression dynamics of representative Gene Ontology (GO) terms across the four stages of EC development. The selected terms reflect biological processes altered during epithelial transformation, including vesicle transport, cytoskeletal regulation, metabolic shifts, and immune-related functions.

Next, we assessed the copy number alteration (CNA) landscape across the major cell types at different EC development timepoints via copyKat^14^, which infers CNAs from transcriptomic data. In epithelial cells, we identified a subgroup exhibiting widespread copy number changes— reflected by higher CNA scores—that became more pronounced during EC progression, a pattern not detected in other cell types (Supplemental Fig. 2b). To characterize genomic instability further, we classified each epithelial cell into three categories on the basis of its genome-wide CNA score, which reflects its CNA burden: genome stable (scores lower than Q3 + IQR), genome moderately stable (scores between Q3 + IQR and Q3 + 3×IQR), and genome unstable (scores at or above Q3 + 3×IQR). The overgrowth (EIN) phase exhibited a significantly greater proportion of moderately stable epithelial cells, whereas the invasive transition and EC groups contained more genome-unstable cells (Fig. 2b). These observations support the notion that early, transient genomic instability evolves into more extensive copy number alterations during the primary tumor expansion phase^15^. We also noted that a greater fraction of epithelial cells lacked detectable CNAs than did tissue samples^7^, likely reflecting our broader sampling of the entire uterine environment rather than the tumor-enriched regions typically sampled in cancer tissues.

### Transcriptomic and functional shifts captured by uterine blood reflect EC development

Using our dataset encompassing four phases of EC development, we applied a generalized linear model with MAST^16^ to identify gene expression changes correlated with disease progression. We calculated expression changes over EC progression and *p*-values for each gene to evaluate their correlation with EC development. Among the 26,546 genes analyzed, 12 were significantly upregulated and 181 were significantly downregulated over the course of EC development (BH adjusted *p*-value < 10^-50^, |log_10_(regression coefficient)| > 0.2) (Fig. 2c). The top DEGs are known to be associated with EC or other cancer types, indicating that uterine blood sampling can reveal biomarkers relevant to cancer progression, similar to tissue samples. Among the top negatively correlated genes, *TRMT9B*—a tRNA methyltransferase gene—has been reported to be homozygously deleted in more than 2% of pancancer samples^17^. Another downregulated gene, *FAM189A2*, encodes a novel ITCH ubiquitin ligase activator previously shown to be significantly reduced in EC tissues and linked to poor prognosis in EC patients^18^. The significantly upregulated genes included *CD24*, a cell adhesion molecule that regulates epithelial‒mesenchymal transition in ovarian cancer^19^ and serves as a poor prognostic marker in EC^20^. Polyadenylate-binding protein cytoplasmic 1 (*PABPC1*) is also elevated; it stabilizes mRNA molecules, increasing protein production, which promotes cancer cell growth, and is dysregulated in ovarian, breast, gastric and hepatocellular carcinomas^21–24^. Additionally, HE4 (*WFDC2*) is known to be overexpressed and promotes tumor growth in EC^25^, whereas *YWHAZ* promotes ovarian cancer metastasis and correlates with unfavorable survival by influencing cancer cell metabolism^26^. Finally, *B2M* plays critical roles in the progression, prognosis, and immunotherapy of gliomas^27^. Gene network analysis^28^ revealed a highly interconnected cluster of genes significantly associated with EC development (Fig. 2d). Within this network, the upregulated genes emerged as hub genes, exhibiting high connectivity and potentially functioning as central regulatory drivers. Functional enrichment analysis of these genes in the Gene Ontology (GO) molecular function domain^29,30^ revealed overrepresented processes such as actin binding (GO:0003779), cell adhesion molecule binding (GO:0050839), growth factor binding (GO:0019838), platelet-derived growth factor binding (GO:0048407), and insulin-like growth factor I binding (GO:0031994) (Fig. 2d). These enriched pathways imply a role in epithelial‒mesenchymal transition (EMT), highlighting the malignant transformation of epithelial cells. Collectively, these findings underscore the capacity of uterine blood analysis to capture key gene expression signatures that drive EC progression.

To gain deeper insight into the functional changes occurring during the transition from a benign endometrium to a malignant endometrium, we identified gene sets correlated with EC development stages via GO modules with MAST^16^. The top 50 correlated GO modules are summarized in Supplemental Figure 2c, and we highlight the module scores for the four most positively and two most negatively correlated modules in Figure 2e, which illustrate gradual functional shifts during EC progression. Notably, we observed downregulation of “Epithelial Cilium Movement Involved In Determination Of Left/Right Asymmetry” (GO:0060287), an intriguing finding given that our analysis focused on unciliated epithelial cells and thus suggested that genes associated with ciliated epithelia may also be altered. A decrease was likewise noted for “Benzene-Containing Compound Metabolic Process” (GO:0042537). Conversely, pathways associated with vesicle transport and exosome assembly, such as “Regulation of Extracellular Exosome Assembly” (GO:1903551), “COPI Coating of Golgi Vesicle” (GO:0048205), and “Golgi Transport Vesicle Coating” (GO:0048200), were upregulated. Additionally, “antigen processing and presentation” pathways presented reduced expression (Supplemental Fig. 2c), potentially reflecting immune evasion by cancerous epithelial cells, whereas “regulation of response to wounding” (GO:1903034) presented increased expression, supporting tumor growth. Other processes, including “Purine Ribonucleoside Monophosphate Catabolic Process” (GO:0009169) and “Release of Cytochrome C from Mitochondria” (GO:0001836), were also implicated in cancer development. These findings underscore the value of liquid biopsy for detecting both gene expression and functional alterations comparable to those found in tissues, highlighting its promise as a liquid biopsy and minimally invasive diagnostic tool for EC.

### Fibroblast lineage remodeling reveals a conserved protumorigenic trajectory

We further investigated fibroblast transcriptome dynamics by performing RNA velocity analysis^13^, which identified the fibroblast lineage trajectory and genes driving the cancer-associated fibroblast phenotype. Velocity-vector mapping revealed unidirectional flow from control fibroblasts through EIN and transition stages toward a distinct EC-specific terminus (Supplementary Fig. 3a). Among the top velocity genes in the EC group, we identified *INHBA*, *ITGA1*, and *CNTN1* (Supplementary Fig. 3b). *INHBA* encodes Activin A, which was recently shown to mark an immunoregulatory CAF subset that induces PD-L1 and promotes T-reg differentiation^31^; its progressive up-shift along the velocity stream is therefore consistent with the increasingly suppressive milieu we documented in immune cells. *ITGA1*, a collagen-binding α1-integrin up-regulated in desmoplastic stroma and several carcinomas^32^, likely equips late-stage fibroblasts for increased matrix engagement and mechanical remodeling. *CNTN1* is an adhesion molecule with pan-cancer pro-invasive functions. We also identified *RBMS3*, an RNA-binding protein implicated in extracellular-matrix regulation and stromal activation^33^, which drives velocity in the EC, transition and EIN stages (Supplementary Fig. 3b), further supporting a model in which velocity-defined fibroblasts acquire both structural and signaling attributes that potentiate tumor invasion. Comparable stromal trajectories—marked by INHBA- or integrin-high CAF subsets—have been reported in pancreas, breast and colorectal cancers^31,34–36^, suggesting that the evolutionary path we detect is part of a conserved, tumor-type-agnostic fibroblast program.

Importantly, these signatures are readily captured in our uterine blood, underscoring the utility of liquid biopsy for identifying expression and functional changes comparable to those detected via tissue and highlighting its potential as a noninvasive diagnostic tool for EC.

### Increased crosstalk between the stroma and uterine microenvironment during endometrial carcinogenesis

In addition to the epithelial compartment, our single-cell atlas encompasses expression profiles from diverse immune populations, providing a comprehensive view of intercellular communication throughout endometrial cancer progression. By mapping ligand–receptor interactions^37^, we observed a consistent increase in the number of signaling events during carcinogenesis and progression (Fig. 3a), with the most pronounced changes arising from ciliated epithelial cells and fibroblasts as senders and myeloid and lymphoid cells as receivers (Fig. 3a, b). These findings align with the concept that developing tumors remodel the microenvironment, intensifying crosstalk and reprogramming immune and stromal cells as malignancy advances^38^.

**Figure 3.**
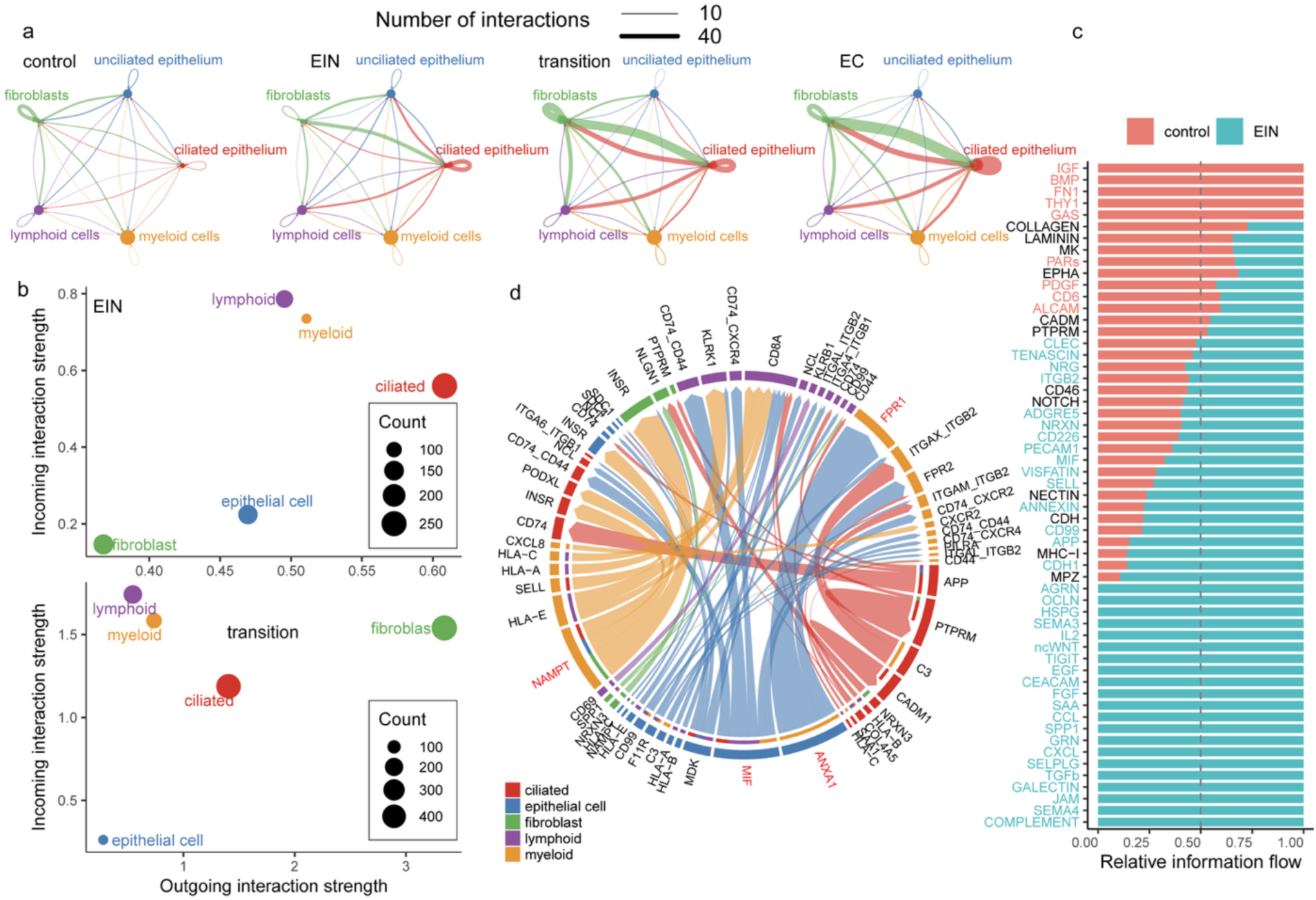
Stromal–epithelial communication intensifies during endometrial tumorigenesis. (a) Network diagrams illustrating the number (edge width) and direction (arrows) of ligand– receptor interactions among major cell classes across the control, EIN and carcinoma stages. (b) Scatterplots showing outgoing versus incoming interaction strength for each cell class (dot size reflects interaction count). (c) Differential information-flow stacked bar plot ranking signaling pathways by the change in total information flow between the EIN and control groups. Significant pathways enriched in the control (red) and EIN (green) groups are highlighted. (d) Chord diagram visualizing ligand–receptor pairs that are significantly up-regulated in EIN versus the control. Ribbons are colored by sender cell type; outer labels indicate ligands and receptors. Pairs discussed in the text are highlighted in red.

When the information flow of signaling pathways from EIN to the control group was compared, multiple growth and adhesion signals—IGF, BMP, FN1, THY1, and GAS—were significantly downregulated (Fig. 3c). In the normal endometrium, these ligands typically help maintain organized tissue architecture and balanced proliferation. For example, IGF supports controlled epithelial growth and can direct macrophages toward an immunoregulatory phenotype^39^; BMP constrains unchecked proliferation^40^; fibronectin shapes the extracellular matrix^41^; THY1 stabilizes fibroblast–epithelial interactions^42^; and GAS6–AXL signaling promotes epithelial survival under homeostatic conditions^43^. Their simultaneous reduction in EIN suggests a breakdown in the usual regulatory crosstalk, permitting nascent neoplastic cells to escape growth constraints and disrupt normal immune oversight. Conversely, the transition to EIN was characterized by upregulated adhesion and immune-associated pathways, such as JAM family proteins, Complement, and SEMA4. These changes reflect an increasingly interactive niche: JAM proteins can both promote tumor cell clustering and alter immune cell trafficking^44,45^; complement activation paradoxically recruits tumor-promoting myeloid cells^46^; and SEMA4D drives myeloid-derived suppressor cell (MDSC) expansion^47^.

A closer look at the upregulated signaling ligand‒receptor pairs revealed that myeloid cells are key participants in these emerging signaling loops (Fig. 3d, yellow). Notable examples include ANXA1–FPR1 signaling from the epithelium to myeloid cells, which can push macrophages toward a tumor-promoting M2 state while suppressing T-cell function^48^, and MIF, a pleiotropic cytokine that orchestrates chronic inflammation yet blunts effective T-cell responses^49^, with increased signals from the epithelium to both myeloid and lymphoid cells. NAMPT signals from myeloid cells to the epithelium and fibroblasts further underscore the metabolic reprogramming of both immune and tumor cells, linking NAD^+^ synthesis to immunosuppression and angiogenesis^50^. The detailed signaling changes in lymphoid and myeloid cells are shown in Supplementary Fig. 4a-d. Collectively, these results underscore the central role of the innate immune system in shaping the tumor microenvironment during EC development.

### Immune cell reprogramming accompanies EC initiation and progression

To quantify how each immune lineage drifts away from its physiological state, we calculated the maximum-mean-discrepancy (MMD) distance^51^ between every cell type at each disease stage and its counterpart in control samples (Supplementary Fig. 5a). We further defined the “speed” of change as the MMD distance between consecutive stages (Supplementary Fig. 5b). Most immune populations reached their greatest transcriptional divergence at the transition stage, underscoring this interval as a critical inflection point. Two exceptions emerged: basophils showed a monotonic decrease in divergence, whereas neutrophils displayed a steady increase. When the change velocity was examined, lymphoid cells (B, T, and NK) maintained a relatively constant pace, whereas most myeloid subsets accelerated markedly throughout disease evolution (Supplementary Fig. 5b), highlighting their dynamic plasticity.

To further investigate myeloid cell remodeling, we first calculated the classical (M1) and alternative (M2) macrophage scores on the basis of the expression levels of curated MSigDB gene sets^52^. We calculated the M1 score on the basis of genes upregulated in macrophages: classical (M1) versus alternative (M2) (GSE5099_CLASSICAL_M1_VS_ALTERNATIVE_M2_MACROPHAGE_UP))^53^ and the M2 score on the basis of (Immunologic Signature (ImmuneSigDB): genes downregulated in macrophages: classical (M1) versus alternative (M2) (GSE5099_CLASSICAL_M1_VS_ALTERNATIVE_M2_MACROPHAGE_DN).^53^ During the control-to-EIN transition, we observed significant changes in accordance with the calculated distance metrics: M1 macrophage signatures significantly decreased, whereas the immunosuppressive M2 signatures increased (Supplementary Fig. 6). This pattern indicates early establishment of an immunosuppressive microenvironment, wherein elevated M2 and reduced M1 macrophages potentially facilitate neoplastic growth.

Collectively, these observations emphasize significant immunological remodeling during the early stages of EC development.

### Neutrophil transcriptomic remodeling during EC invasive transformation

Among all the cell types examined, neutrophils exhibited the most pronounced and statistically significant alterations throughout EC progression. Continuous increases in transcriptomic divergence from the control group were observed with increasing disease stage (Supplementary Fig. 5a). This dynamic pattern was further substantiated by the spatial distribution of neutrophil populations visualized through kernel-density estimation contours projected onto a UMAP embedding (Fig. 4a). Each contour represents neutrophil density distributions corresponding to distinct pathological stages (control, EIN, transition, and EC), with arrows linking the centroid positions of each population to illustrate directional progression. A clear trajectory emerged from the control state toward the EC state, with the most substantial shift occurring between the EIN state and the transition state—where endometrial cells begin acquiring invasive characteristics.

**Figure 4.**
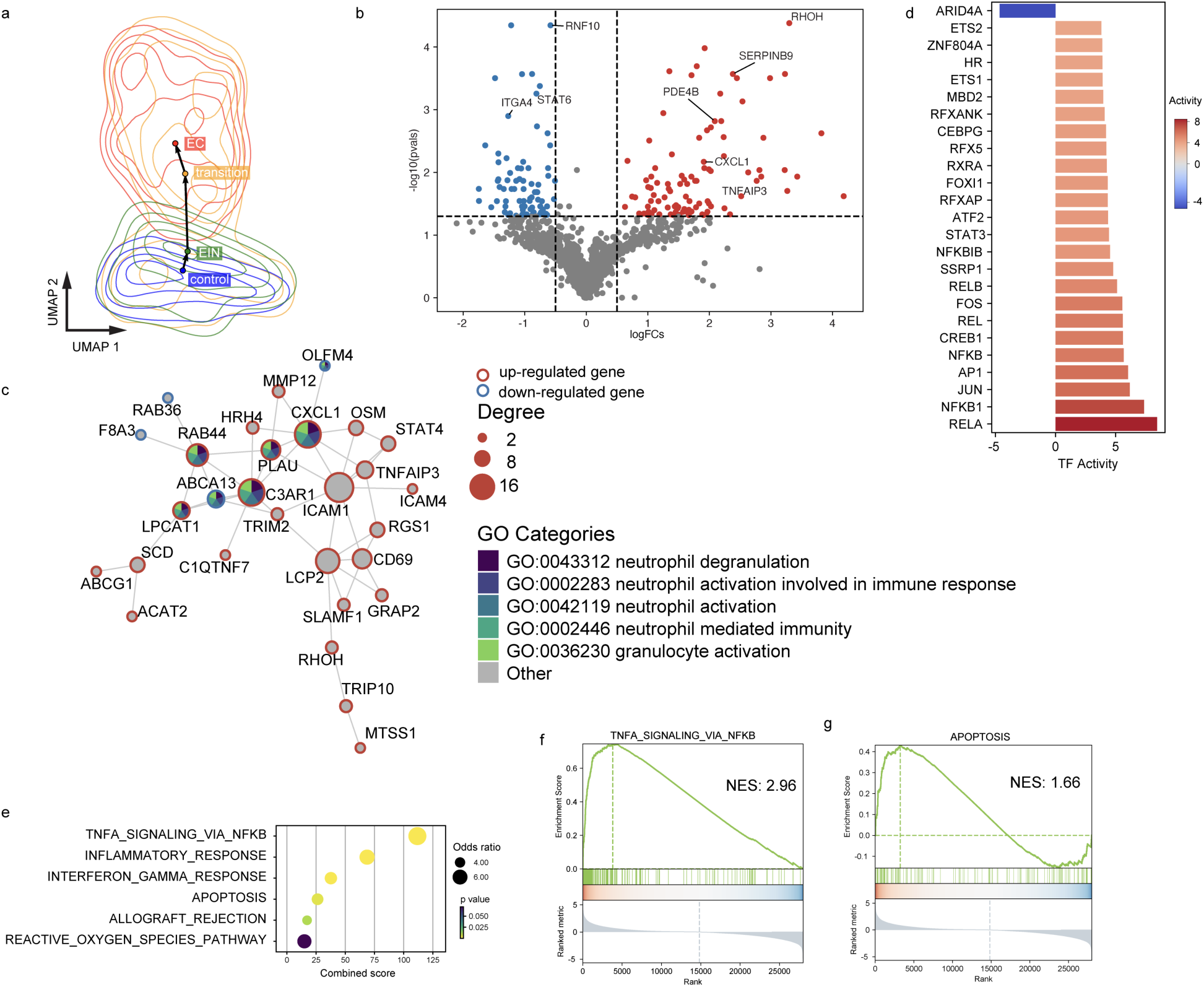
Neutrophil re-programming during the invasive transformation of ECs. (a) Kernel-density contours of neutrophils projected onto a shared UMAP; arrows connect centroid positions across stages. (b) Volcano plot of differentially expressed genes (DEGs) (invasive vs non-invasive neutrophils); selected genes are labeled. (c) STRING protein–protein interaction network of significant DEGs; node size reflects degree, color denotes enriched GO terms related to neutrophil function, and outline color denotes the direction of regulation in invasive versus non-invasive neutrophils (red = up-regulated, blue = down-regulated). (d) Top transcription factors ranked by activity change; bars to the right (red) are activated in invasive neutrophils, and bars to the left (blue) are repressed. (e) Over-representation analysis for hallmark gene sets; dot size = odds ratio, color= *p*-value. (f–g), GSEA plots showing positive enrichment of TNFα signaling via NF-κB (f) and mixed enrichment of apoptosis genes (g) in invasive neutrophils. NES, normalized enrichment score.

To identify key molecular changes during critical invasive transition, we performed differential gene expression analysis on neutrophils^54^, combining control and EIN as the noninvasive category and transition plus EC as the invasive category. By comparing invasive and noninvasive neutrophil populations, we identified 98 significantly upregulated genes and 77 significantly downregulated genes (|log₂FC| > 0.5, q value < 0.05; Fig. 4b). Functionally, the transcriptomic shift that accompanies invasive transformation indicates that neutrophils are rewired to a highly proinflammatory state while equipping them with mechanisms to evade immune attack. Notably, several negative regulators of inflammation or motility are repressed in invasive neutrophils. For example, *RNF10,* an E3 ubiquitin ligase that terminates NF-κB/IRF3 signaling, is downregulated, which is consistent with relieving a brake on inflammatory pathways^55^. *STAT6* expression is also reduced, which is in line with reports that STAT6 normally restrains neutrophil-driven inflammation^56^. Similarly, *ITGA4* (integrin α4), which helps anchor myeloid cells in stromal niches^57^, is diminished, potentially enabling greater neutrophil mobility and tissue infiltration. The loss of these restraining factors is coherent with a switch toward an activated, cytokine-secreting neutrophil phenotype that is both highly migratory and immunosuppressive.

Conversely, neutrophils in the invasive stage upregulate a suite of effector genes that amplify inflammation, increase survival, and promote the recruitment of additional myeloid cells. For example, *PDE4B* is significantly induced. PDE4B lowers the level of intracellular cAMP, thereby unleashing respiratory-burst activity and chemotaxis^58^; PDE4B has a proinflammatory role and has been shown to facilitate inflammation-driven tumorigenesis^59^. In parallel, invasive neutrophils show increased autocrine production of *CXCL1*, a chemokine that establishes a feed-forward loop to attract additional CXCR2⁺ neutrophils and myeloid cells into the tumor invasive front^60,61^.

Additionally, the cytotoxic resistance of invasive neutrophils is reinforced via the upregulation of *SERPINB9* (also called PI-9), an inhibitor of granzyme B that protects cells from granzyme-mediated apoptosis^62^. Elevated *TNFAIP3* (A20) further insulates these neutrophils by providing a negative feedback brake on NF-κB signaling, preventing runaway inflammatory activation that could trigger cell death^63^. Finally, *RHOH* is strongly induced; RHOH is a unique Rho GTPase that dampens Rac-dependent actin dynamics^64^. Increased RHOH serves to restrain excessive degranulation and NETosis^65^, thereby limiting collateral tissue damage while still preserving tumor-promoting inflammation. Taken together, these findings indicate that the invasive transition is marked by the loss of key anti-inflammatory regulators and the acquisition of proinflammatory, prosurvival programs by neutrophils, resulting in a specialized immunosuppressive neutrophil phenotype. This reprogrammed neutrophil state appears to drive the leap from EIN to the transition stage and sustains the invasive progression of EC by simultaneously promoting inflammation and protecting the tumor microenvironment from immune elimination.

### Functional network analysis reveals convergent protumor signatures

To explore the functional relationships among these differentially expressed genes (DEGs), we constructed a protein‒protein interaction network (Fig. 4c) via the STRING database^28^. In this network, nodes represent the proteins encoded by significant DEGs, and edges represent known or predicted interactions. We annotated each node with the top 5 Gene Ontology Biological Process (GO BP) terms^29,30^ overrepresented among the DEGs. In the network, the hub genes that are highly connected and enriched in functions related to neutrophil activation and immune effector processes, such as “neutrophil degranulation”, “neutrophil activation”, “neutrophil-mediated immunity” and “granulocyte activation”, are involved. The highly connected hub genes that were upregulated included *CXCL1*, *C3AR1* (the receptor for C3a), *RAB44* (involved in granulocyte exocytosis), and *LPCAT1* (linked to inflammatory lipid mediator production). These hub genes indicate enhanced secretory and signaling functions in invasive neutrophils. In contrast, several downregulated DEGs were associated with the negative regulation of neutrophil activation and maturation. For example, *OLFM4*, a gene typically expressed in a subset of neutrophils with more suppressive or immature characteristics, is significantly diminished^66^. The loss of OLFM4 and similar genes could indicate a shift away from a regulatory neutrophil subset toward a fully activated subset. Overall, this network analysis underscores the dramatic functional shift in neutrophils during the invasive transition: the neutrophil DEGs are not random but instead converge on pathways of cell activation, secretion, motility, and immune defense—all hallmarks of an “aggressive” neutrophil phenotype geared to support tumor invasion.

### Transcriptional control of invasive neutrophil identity

We next sought to identify the upstream transcription factors (TFs) that might drive this neutrophil reprogramming. Using the CollecTRI framework^67^, a comprehensive resource that integrates TF-target interactions from 12 databases, we computed the activity of transcriptional regulators in invasive vs. noninvasive neutrophils. CollecTRI considers not only the differential expression of TFs but also the coordinated up-/downregulation of their known target genes, providing a robust inference of TF regulatory activity (with directionality, i.e., activation or repression). The analysis revealed a set of master regulators whose activities shifted significantly during the neutrophil invasive transition (Fig. 4d lists the top 15). Among these factors, NF-κB and AP-1 family factors have emerged as dominant positive regulators in invasive neutrophils. In particular, RELA (NF-κB p65 subunit) and NFKB1 (p50 subunit) showed substantially increased activity, highlighting that canonical NF-κB signaling is highly active in neutrophils at the invasive stage, which is consistent with the inflammatory milieu of the tumor microenvironment^68^. Concurrently, JUN, a component of the AP-1 complex (c-Jun), was upregulated and more active. AP-1 (often a dimer of Jun/Fos proteins) is commonly coactivated alongside NF-κB in response to proinflammatory cytokines and stress, and it cooperates with NF-κB to induce the expression of many cytokine and chemokine genes^69–71^. The enrichment of AP-1 and NF-κB suggests a synergistic transcriptional program promoting inflammation, survival, and tissue remodeling functions in neutrophils.

In contrast, the analysis indicated decreased activity of certain transcriptional repressors in invasive neutrophils. Notably, ARID4A (AT-rich interactive domain 4A, also known as RBBP1) was identified as a significantly downregulated regulator. ARID4A is a known corepressor that can be recruited by the retinoblastoma protein (RB1) to inhibit E2F target genes^72^. Its reduced activity in invasive neutrophils aligns with the derepression of proliferation- or cell cycle-related genes. Indeed, our TF-target network (Supplementary Fig. 7a) revealed that ARID4A normally represses certain genes (e.g., cell cycle regulators such as *RB/E2F* targets), but other upregulated TFs (AP-1, NF-κB, etc.) drive those same genes in the opposite direction. For example, ARID4A has been reported to repress RB1 and other cell cycle checkpoint genes, but in invasive neutrophils, RB1 itself and many E2F-regulated genes trend upward, likely because the combined activation of AP-1 and NF-κB overcomes the loss of ARID4A repression. Another illustrative target is CXCL1: this chemokine is among the top upregulated genes and is a shared target of multiple activated TFs (NF-κB, AP-1, etc.), all of which are consistently proinflammatory regulators. The concordant upregulation of both the TFs and their proinflammatory targets (such as *CXCL1*, *IL1B*, and TNFAIP3) indicates a robust, self-reinforcing regulatory network. In summary, the invasive neutrophil state is orchestrated by a shift in the balance of transcriptional regulators: proinflammatory transcription factors (NF-κB, AP-1, etc.) are intensified, whereas anti-inflammatory or differentiation-related repressors (such as ARID4A) are attenuated. This creates a synergistic regulatory circuit that locks neutrophils into a tumor-promoting activation state.

### Invasive neutrophils are enriched in hallmark inflammatory and survival pathways

To confirm the functional implications identified from these gene expression changes, we performed an overrepresentation analysis (ORA)^54^ of the MSigDB hallmark gene sets^52^ using the DEGs. The ORA results (Fig. 4e) highlighted several pathways that were significantly enriched in the neutrophil invasive signature. The top enriched terms identified were TNFα signaling via NF-κB, the inflammatory response, the interferon gamma response, apoptosis, allograft rejection, and the reactive oxygen species (ROS) pathway. These enriched terms strongly reinforce the notion that invasive neutrophils undergo immunological and metabolic metamorphosis. The identified pathways underscore the dual role of the inflammatory response in cancer progression. On the one hand, inflammation can trigger immune-mediated destruction of abnormal cells, but on the other hand, chronic inflammation and associated signaling pathways—such as TNFα signaling via NFκB—can promote tumor progression by increasing cellular proliferation, survival, and metastasis^73^. The enrichment of pathways such as the interferon gamma response highlights the complex interplay between the immune system and the tumor microenvironment^74^. These pathways may reflect an ongoing but ineffective immune response, where tumor cells evade immune surveillance while exploiting immune signals to promote their growth. The involvement of wound-healing pathways, such as the inflammatory response and allograft rejection, suggests a shift in the microenvironment toward tissue remodeling and repair processes^75^. This is consistent with the creation of a “progrowth” niche that facilitates tumor invasion and metastasis.

### Neutrophils fine-tune apoptosis and the ROS balance during invasion

Perhaps the most intriguing findings are the enrichment of apoptosis and ROS pathway hallmarks, given the complex role of neutrophils in cell death and survival. In terms of facial value, one might expect prosurvival changes to allow neutrophils to persist in tumors and possibly a dampened oxidative burst to avoid tissue damage that could hinder tumor growth. Our data indeed suggest that invasive neutrophils adopt a more survival-prone, antiapoptotic profile (e.g., upregulating SERPINB9 and A20, as discussed previously) and modulate their oxidative machinery (e.g., upregulating RHOH). The enrichment of an apoptosis gene set could seem paradoxical—why would a cell type that we propose to be more resistant to death show elevated expression of apoptosis-related genes? The likely explanation is that this hallmark gene set contains both proapoptotic and antiapoptotic genes, and neutrophils in the invasive stage tweak both sides of that equation.

To further dissect the seemingly paradoxical enrichment of both the ROS-related and apoptosis-related signatures, we performed a more granular gene set analysis focusing on the enriched gene sets. Gene set enrichment analysis (GSEA)^76^ was applied to the hallmark terms of interest (Fig. 4f, g; Supplementary Fig. 7b–e). This analysis revealed that certain hallmarks, such as TNFα signaling, were positively enriched predominantly due to upregulated genes (Fig. 4f), whereas others, such as apoptosis (Figure 4g) and the reactive oxygen species pathway (Supplementary Fig. 7c), showed split enrichment: subsets of those gene sets were enriched among upregulated genes and other subsets among downregulated genes. In other words, invasive neutrophils upregulate some apoptosis-related genes while repressing others, similar to ROS-related genes. This complex rebalancing suggests that the shift from noninvasive to invasive neutrophils is not a uniform on/off switch for ROS or apoptotic activity but rather a fine-tuning of these programs.

### Rebalancing of the ROS and apoptosis programs in invasive neutrophils

We hypothesized that this phenomenon reflects a switch from a pro-oxidant, pro-apoptotic state in noninvasive neutrophils to an antioxidant, anti-apoptotic state in invasive neutrophils. To test this, we calculated module scores for curated gene panels (derived from prior studies of neutrophils in gliomas^77^) that classify genes as either promoting or counteracting ROS and apoptosis. Indeed, the results (Fig. 5a-c; Supplementary Fig. 8) demonstrated that noninvasive neutrophils (control/EIN) presented significantly increased expression of pro-ROS and proapoptotic genes, whereas invasive neutrophils (transition/EC) presented elevated anti-ROS and antiapoptotic signature scores.

**Figure 5.**
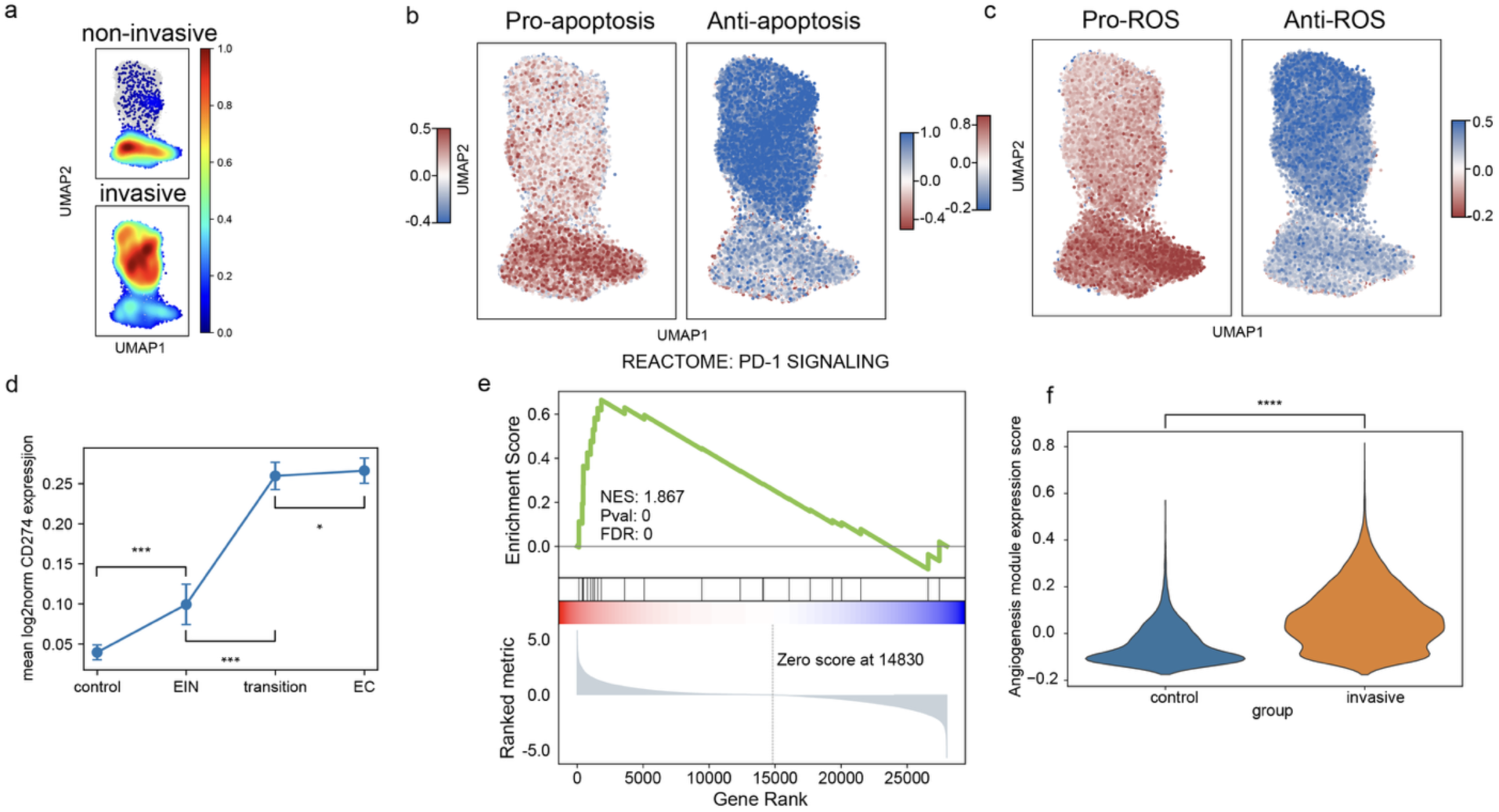
Functional re-wiring of neutrophils during the invasive transition. (a) Kernel-density contours (UMAP) highlighting the spatial redistribution of neutrophils from non-invasive to invasive stages. (b–c) UMAPs depicting the balance between opposing gene programs in individual neutrophils. (b) Apoptosis module: Red cells express a predominance of pro-apoptotic genes, whereas blue cells are enriched for anti-apoptotic genes. (c) ROS module: Red cells favor pro-ROS genes, whereas blue cells favor anti-ROS genes. (d) Stage-wise increase in mean log-normalized *CD274* (PD-L1) expression (mean ± s.e.m.; Mann-Whitney). (e) GSEA plot demonstrating positive enrichment of the REACTOME PD-1 signaling gene set in invasive neutrophils. (f) Violin plots of angiogenesis-module scores in control/EIN versus transition/EC neutrophils (\*\*\**P* < 0.0001, Mann-Whitney test).

This finding aligns well with observations in other tumor contexts^77^; for example, tumor-associated neutrophils in brain tumors downregulate NADPH oxidase components (such as NCF1, NCF2, and NCF4) while upregulating antioxidant enzymes (such as the glutathione-synthesis enzymes GCLC and GCLM and the peroxiredoxins PRDX1/2/4)^77^. Functionally, this means that invasive neutrophils produce a more controlled, lower level of ROS enough to signal and modulate the environment but not so much as to cause excessive tissue damage or induce their own cell death. Similarly, we found that invasive neutrophils in ECs suppress many proapoptotic factors and instead induce the expression of survival genes, including known apoptosis inhibitors such as IAPs (BIRC2 (cIAP1) and BIRC3 (cIAP2))^77^ and antioxidative stress proteins. This shifts the balance toward an oxidative stress-resistant, apoptosis-resistant state.

The reprogramming of neutrophils at the invasive stage is biologically sensible in the context of tumor progression. Early-disease-state neutrophils (e.g., in EIN lesions) may produce bursts of ROS as part of the normal immune response to emerging cells, potentially exerting initial antitumor effects. They may also be more prone to activation-induced cell death after performing their immune functions. However, as the tumor continues to grow and coopts the microenvironment, the neutrophils that persist and infiltrate the invasive front are those that have adapted to a protumor phenotype. This includes reducing collateral damage from ROS and avoiding cell death. By curtailing ROS production (through lower NADPH oxidase activity and higher antioxidant expression), these neutrophils minimize oxidative damage to surrounding tumor cells and tissues, preventing an excessive inflammatory burst that could harm tumor expansion. Moreover, by upregulating antiapoptotic mechanisms, neutrophils themselves survive longer in the TME. Indeed, invasive tumor-associated neutrophils are known to prolong lifespan and immunosuppressive capacity^77^, behaving more like chronic inflammatory cells than short-lived first responders do.

### Invasive neutrophils acquire immunosuppressive and proangiogenic phenotypes

Neutrophils in early lesions presented minimal PD-L1 (*CD274*) level, but *CD274* expression levels increased progressively across the control, EIN, transition, and invasive carcinoma samples, reaching the highest expression in ECs (Fig. 5d). Each developmental transition was associated with a significant increase in neutrophil PD-L1 (*p* < 0.05), indicating that during the invasive stage, neutrophils acquire an immune checkpoint ligand highly associated with T-cell suppression. Emerging evidence indicates that tumor-associated neutrophils (TANs) can upregulate the expression of the immune checkpoint ligand PD-L1 as cancers progress, contributing to an immunosuppressive TME. In several solid tumors, infiltrating neutrophils exhibit high surface PD-L1 expression, which is correlated with advanced disease stage and poor patient outcomes^78,79^. In parallel, gene set enrichment analysis revealed that invasive-stage neutrophils were significantly enriched for the REACTOME^80^ : PD-1 signaling gene set (normalized enrichment score 1.867, FDR ≈ 0; Fig. 5e). This enrichment of the PD-1/PD-L1 checkpoint pathway, together with the upregulation of PD-L1 itself, suggests that neutrophils in the tumor microenvironment actively engage in PD-1–mediated immune evasion. Indeed, neutrophils expressing PD-L1 are known to directly inhibit cytotoxic T cells and promote immune escape in other cancers^77^. Consistent with this immunosuppressive TAN phenotype, our data implicate the PD-1/PD-L1 axis as a key mechanism by which neutrophils facilitate immune suppression in invasive ECs.

In addition to immune modulation, invasive neutrophils adopt a pronounced proangiogenic program. A curated 20-gene angiogenesis signature was markedly upregulated in neutrophils from invasive tumors, including canonical growth-factors such as *VEGFA* and *FGFR1*, ECM restructuring versican (*VCAN*) and periostin (*POSTN*) as described in other tumor contexts^77^, with module scores significantly higher than those in noninvasive samples (Mann–Whitney test, *p*<0.0001; Fig. 5f). This result indicates that as endometrial lesions become invasive, neutrophils begin actively supporting tumor vascularization. Tumor-associated neutrophils have been shown to secrete potent proangiogenic factors, including VEGF (also validated in our data with increased VEGFA), which remodel the extracellular matrix and stimulate new blood vessel growth^81^. The elevated angiogenesis score in invasive-stage neutrophils aligns with these reports, reflecting an acquired proangiogenic phenotype likely to enhance the blood supply to the tumor.

### Temporal transcriptome dynamics reveal neutrophil progressive adaptation to the tumor microenvironment

We then calculated the genes whose expression patterns consistently increased or decreased across the developmental stages with TDEseq^82^ for genes whose temporal gene expression patterns increased during EC development. We defined genes that rose continuously from control through EIN, transition, to carcinoma (“growth” genes) (Fig. 6a, red cluster). Functional enrichment analysis revealed that these genes were enriched in immune activation pathways, including the NF-κB pathway, cytokine-mediated signaling and apoptotic signaling (Fig. 6b). Our previous noninvasive vs invasive analysis revealed gradual changes across the stages of EC development. Neutrophils increasingly engage in proinflammatory transcriptional programs and activation, which is consistent with neutrophils responding to chronic inflammatory cues in the tumor microenvironment, prolong neutrophil survival and skew them toward tumor-associated phenotypes.

**Figure 6.**
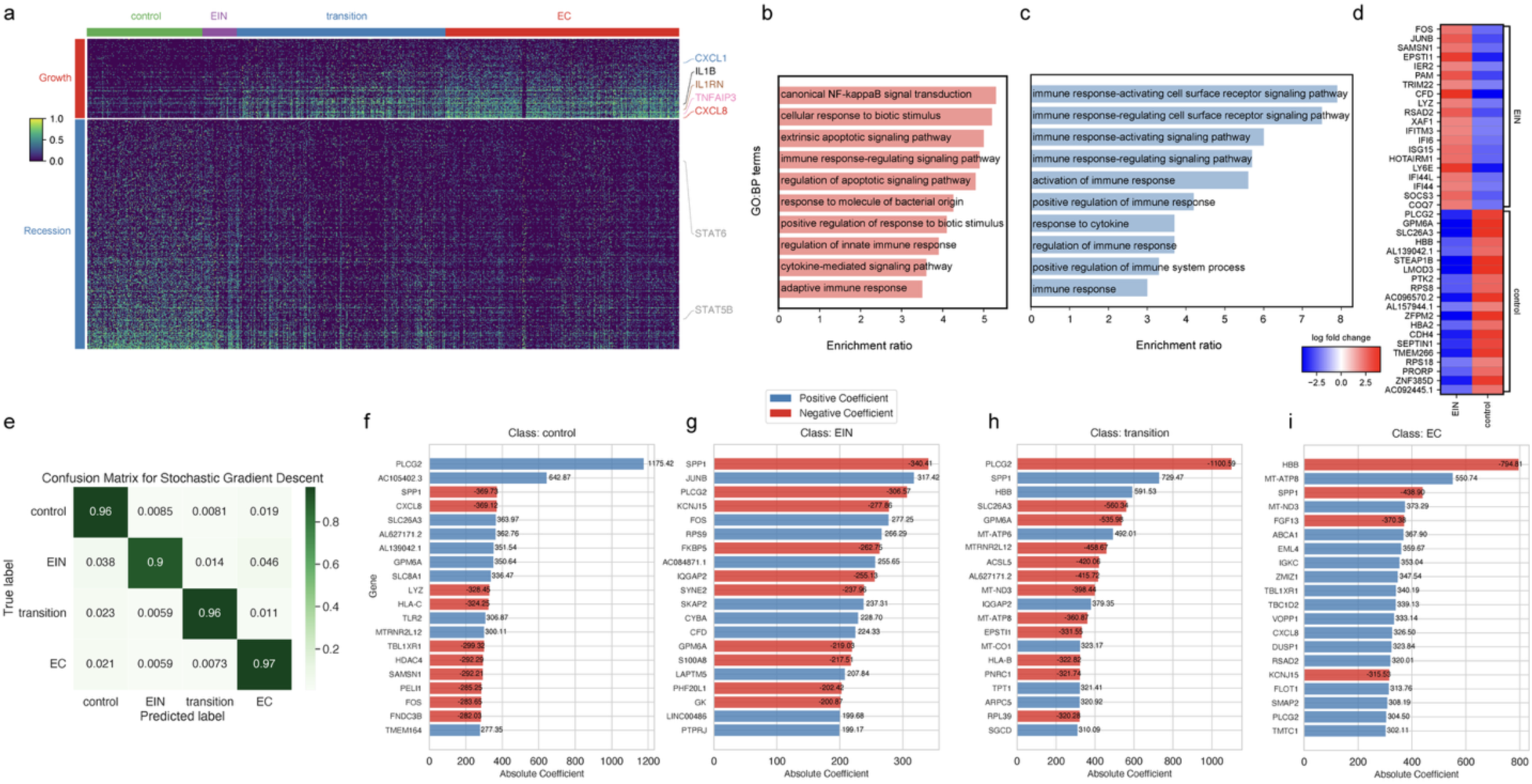
Gene expression and functional changes in neutrophils during EC progression. (a) Heatmap of stage-ordered neutrophil transcriptomes (rows = cells, columns = genes). Genes were clustered into “growth” (red) and “recession” (blue) modules by TDEseq; the color bar indicates z-scored expression. (b–c) GO-BP enrichment for the growth (b, salmon) and recession (c, blue) modules; bars show the enrichment ratios. (d) Heatmap of log₂-fold changes for significantly deregulated genes in the EIN group versus the control group; red = up, blue = down. (e) Confusion matrix of the SGD classifier (values = proportion of true labels). (f–i) Top positive (blue) and negative (red) model coefficients for genes driving the classification of control (f), EIN (g), transition (h) and EC (i) neutrophils; the bar length denotes the absolute coefficient.

In contrast, genes that were monotonically downregulated across these stages (“recession” genes; Fig. 6a, blue cluster) were enriched for broader immune response pathways, such as immune response–regulating receptor signaling and cytokine response (Fig. 6c). This pattern implies that neutrophils gradually attenuate certain general receptor-mediated signaling programs as ECs progress. In the early stages, neutrophils may express a broad array of receptors for immune surveillance, but during tumor development, they possibly downregulate some innate signaling pathways, which is consistent with a shift toward a specialized tumor-associated neutrophil state. Such a recession of broad responsiveness might reflect adaptation or even a form of “tolerance” to the persistent inflammatory environment. The net result is a transcriptional bifurcation: neutrophils in advanced lesions upregulate specific inflammatory and survival pathways while repressing some baseline immune functions, potentially to avoid overactivation or exhaustion in the chronic tumor microenvironment. A complementary correlation analysis further underscored neutrophil adaptation to the tumor environment (Supplementary Fig. 9a, b). This approach identified GO biological processes^29,30^ whose expression in neutrophils was highly correlated with disease stage. Several processes are positively correlated with progression: neutrophil extravasation, interleukin-23 (IL-23) production, T-cell polarization, response to monoamines, and transcriptional regulation in hypoxia. The prominence of extravasation genes in later stages suggests that neutrophils increasingly express programs for tissue infiltration as tumors develop.

### Neutrophils are transcriptionally primed in early EIN lesions

We then focused on the early-stage changes in ECs by comparing the control stage to the EIN stage to identify early changes in neutrophils. Differential expression analysis (Wilcoxon rank-sum test on log-normalized counts) of neutrophils from EIN lesions versus controls revealed robust transcriptomic alterations (Fig. 6d)^83^. EIN-associated neutrophils showed significant upregulation of a subset of immediate-early and inflammatory genes (logFC > 2), including *FOS* and *JUNB* (components of the AP-1 transcription factor complex), and interferon-stimulated genes (*IFITM3*, *ISG15*, *IFI44L*) (Fig. 6d). The elevated expression of *FOS* and *JUNB* is indicative of early neutrophil activation at the EIN, as AP-1 family members are rapidly induced by proinflammatory stimuli^84^. Concurrent upregulation of canonical ISGs such as *IFITM3*, *ISG15*, and *IFI44L* suggests the engagement of type I interferon signaling in EIN neutrophils^85^.

The functional enrichment of these upregulated genes underscores the immune-activated phenotype of EIN neutrophils (Supplementary Fig. 9c). The top enriched Gene Ontology terms centered on cytokine signaling and secretion, including “response to cytokine” and “regulation of cytokine production,” as well as enrichment in neutrophil granule-related components such as the “secretory granule membrane.” Pathway analysis further highlighted interferon α/β signaling (Reactome^80^) and the TNF signaling pathway (KEGG^86^) among the most overrepresented pathways, which is consistent with a proinflammatory and type I interferon-rich milieu in EIN. Together, these transcriptional changes indicate that neutrophils are among the first immune cells to sense and respond to nascent tumor-associated inflammation in EIN lesions. EIN neutrophils thus appear in an “alert” or primed state – marked by AP-1 activation, interferon-response gene expression, and cytokine feedback induction – suggesting early engagement in immunomodulatory or even tumor-promoting functions within the incipient tumor niche. Notably, this early neutrophil priming likely lays the foundation for their later reprogramming into immunosuppressive, tumor-supporting phenotypes as the disease progresses through the transition stage to established endometrial carcinoma.

### Neutrophil transcriptomes predict EC developmental states and reveal key genes

Since such significant changes occur during different stages of EC development, we built a model to predict an individual’s EC developmental stage solely from the transcriptomic profile of their neutrophils. We trained a machine learning model with a stochastic gradient descent model to predict the development stages from which the neutrophil originates on the basis of the transcriptome. We achieved excellent stage classification performance. The model accurately identified the developmental stage of individual neutrophil profiles with minimal confusion between classes, as evidenced by the nearly diagonal confusion matrix (Fig. 6e). Each stage was predicted with ∼90–97% accuracy, and importantly, fewer than 5% of cells were misassigned across the noninvasive (control/EIN) versus invasive (transition/EC) boundary, underscoring the classifier’s strong ability to distinguish early lesions from invasive cancer.

Interrogating the sparse model coefficients revealed distinct gene expression signatures for neutrophils at each stage (Figure 6f-i). For example, JUNB and FOS (AP-1 transcription factor components) had high positive coefficients in classifying EIN-stage neutrophils. The notably high expression of hemoglobin-β (HBB) — known to be induced in activated myeloid cells to help neutralize reactive oxygen species^87^ — coupled with a negative coefficient for the epithelial ion transporter SLC26A3 suggest that neutrophils shift toward an oxidative stress-adapted phenotype as they infiltrate the invasive front. In contrast, EC-stage neutrophils were distinguished by a very large *negative* coefficient for HBB and high positive coefficients for the chemokine CXCL8 (IL-8), along with elevated DUSP1 (a MAPK phosphatase that restrains proinflammatory signaling)^88^ and a negative coefficient for KCNJ15. This pattern implies an immunosuppressive, protumoral activation state for neutrophils in established carcinoma lesions. The concurrent increase in IGKC and ABCA1 in EC-stage neutrophils further hints at crosstalk with B-cell/antibody complexes and altered lipid handling, respectively—both traits observed in tumor-associated neutrophils. In fact, TANs have been shown to interact with and recruit B cells to the tumor microenvironment^89^, where they can accumulate tumor-derived lipids that reprogram them from an antitumor phenotype to a tumor-supporting phenotype^90^. Collectively, these gene expression changes indicate that as neutrophils progress from preinvasive to invasive tumor stages, they transition from an oxidative stress-coping early phenotype to a more immunoregulatory, tumor-supporting phenotype, mirroring the known plasticity of tumor-associated neutrophils in cancer.

These stage-specific neutrophil gene signatures not only provide mechanistic insight into how neutrophil phenotypes evolve from precancer to invasive cancer but could also be leveraged for noninvasive diagnostics. For example, one could envision transcriptomic profiling of neutrophils from easily accessible samples such as menstrual blood to infer an individual’s endometrial cancer stage. In summary, this machine learning approach has the capacity to predict a patient’s disease stage from their neutrophil transcriptional signature, offering both a window into the immune dynamics of endometrial cancer progression and a promising translational tool for early detection and monitoring.

## Discussion

In this study, we present a single-cell atlas of the endometrial cancer microenvironment obtained from uterine blood samples, highlighting profound immune and stromal remodeling even at the precancerous stage. Our key finding is that coordinated reprogramming of immune cells and stromal‒epithelial interactions begins at the EIN stage, well before carcinoma develops. These findings underscore the concept that tumor microenvironment dynamics are integral to early tumorigenesis.

Methodologically, our approach is unique in leveraging uterine cavity blood as a form of local liquid biopsy for high-resolution cellular analysis. We applied droplet-based scRNA-seq directly to uterine blood without enrichment, thereby sampling a wide array of cell types, including endometrial cells, immune cells, and stromal cells, in an unbiased manner. The success of this approach is that uterine blood can serve as a rich source of information about the tumor and its microenvironment, providing a less invasive and more sensitive complement to tissue biopsy. By integrating diverse computational tools, we ensured a comprehensive analysis of cellular states and interactions across disease stages.

Our results provide several important insights. First, immune cell reprogramming is initiated extremely early in the course of EC development. Leukocytes from EIN lesions exhibit significant compositional and transcriptional changes compared with those from cancer-free controls. For example, innate immune cells such as neutrophils display transcriptomic changes suggestive of altered activation states as early as the EIN state. These findings support a model in which the immune microenvironment is actively remodeled in response to early neoplastic transformation. Notably, we demonstrate that neutrophil transcriptomes alone are sufficient to accurately predict an individual’s EC developmental stage, highlighting their potential as noninvasive biomarkers. Moreover, uterine blood captures epithelial and stromal changes that mirror those observed in tissue biopsies, enabling a more complete and representative molecular snapshot of the uterine landscape.

Importantly, our findings also reflect and address key clinical challenges in EC diagnosis. Retrospective analyses have shown that up to 40% of women diagnosed with EIN already harbor concurrent, undetected carcinoma at the time of hysterectomy^91^. This high false-negative rate stems in part from the limitations of endometrial biopsy, which samples only a small, often nonrepresentative portion of the endometrial lining. As a consequence, many patients with EIN undergo a precautionary total hysterectomy even when no invasive cancer is present, leading to overtreatment in a substantial subset of cases. Our uterine blood-based approach offers a compelling alternative: by sampling the uterine environment, we can detect not only epithelial transformation but also immune cell alterations that accompany early tumor evolution. This broader window into disease biology enhances diagnostic sensitivity and provides a molecular framework for understanding how immune remodeling contributes to progression.

Finally, our work identified transcriptional signatures and regulatory networks that are stage-specific and functionally relevant to EC pathogenesis. From the activation of inflammatory and immune checkpoint programs to the emergence of proangiogenic and immunosuppressive neutrophil phenotypes, we reveal key molecular pathways and cell–cell interactions that drive the transition from EIN to invasive carcinoma. These insights identify promising biomarker candidates for early detection and risk stratification and may also inform future strategies for conservative clinical management in patients with premalignant lesions.

Our findings establish a proof-of-concept that cellular and molecular features recovered from uterine cavity blood can serve as the basis for EC diagnostics. Although our specimens were collected intraoperatively, they recapitulate the epithelial, stromal, and immune compartments present in menstrual effluent, supporting the translational premise that naturally shed menstrual blood could be used to detect premalignant and malignant changes. This approach offers advantages over endometrial biopsy, including reduced sampling, feasibility for longitudinal monitoring, and improved patient acceptability, while capturing both tumor-intrinsic and microenvironmental signals that are difficult to measure in small tissue cores.

Critically, these data motivate prospective validation using true menstrual effluent collected with standardized pads/cups and stabilization media. A locked, clinically deployable assay— whether targeted transcript panels or alternative readouts derived from our single-cell signatures—should be evaluated for sensitivity/specificity to (i) distinguish benign vs. EIN vs. EC, (ii) triage EIN for conservative management or definitive surgery, and (iii) enable post-treatment surveillance. Ultimately, head-to-head blinded studies comparing biopsy alone vs. biopsy with liquid biopsy—or non-inferiority designs comparing biopsy to a menstrual-blood test—will be required to define clinical utility and pathways to adoption.

Together, this study advances the mechanistic understanding of endometrial cancer initiation and progression and highlights the value of uterine blood as a high-resolution, minimally invasive diagnostic medium. By capturing the cellular and molecular coevolution of tumor and immune compartments, our approach opens new avenues for both basic discovery and translational application in gynecologic oncology.

## Methods

### Sample collection and processing

Sample collection for this study was approved by the Institutional Review Board at Creighton University in Omaha, Nebraska. Eligible patients were identified prospectively upon chart review by the study team with preoperative endometrial sampling indicating benign, EIN, or EC pathology. Sample collection was performed at the time of planned hysterectomy. Patient data was abstracted through chart review. Data collected included patient demographics, preoperative pathologic diagnosis and endometrial sampling method, and final pathology diagnosis and surgical procedure.

Uterine blood samples were obtained from 15 individuals representing four states of endometrial cancer (EC) development: cancer-free, endometrial intraepithelial neoplasia (EIN), a transitional state of both EIN and EC concurrently present, and pure EC. At the time of planned hysterectomy, an endometrial pipelle was used to aspirate blood from the endometrial cavity. Each sample was processed fresh within 1 hour of collection to preserve RNA integrity. Red blood cells were removed by lysis, and the remaining nucleated cells were washed and resuspended in PBS with 0.04% BSA. No prior cell enrichment or cryopreservation was performed, ensuring an unbiased single-cell suspension directly from the uterine microenvironment.

### Single-cell RNA sequencing

Single-cell RNA sequencing libraries were prepared via the 10x Genomics Chromium platform and 3ʹ Gene Expression v4 chemistry. In brief, single-cell suspensions with a concentration of 1000 cells/μL were loaded onto a Chromium microfluidic chip, where the cells were encapsulated into nanoliter oil droplets (GEMs) together with barcoded gel beads and reagents. Reverse transcription and cell barcoding were performed within each droplet, after which the cDNA was pooled, amplified, and used to construct sequencing libraries following the manufacturer’s protocol. Libraries from all 15 samples were sequenced on an Illumina NextSeq 2000 sequencer to an adequate depth to capture diverse transcriptomes. The raw base call files were demultiplexed and converted to FASTQ format via the Cell Ranger toolkit (10x Genomics).

### Preprocessing and quality control

The raw sequencing reads were aligned to the human genome (GRCh38) via Cell Ranger v7.1.0^92^, which generated gene‒cell count matrices for each sample. Downstream analysis was performed in Python via the Scanpy library (v1.9)^83^ for single-cell data processing. We first filtered out empty droplets and low-quality cells. To detect and remove doublets, we employed Scrublet^93^, which simulates doublets from the dataset and scores each cell for doublet likelihood. Cells with high Scrublet scores were flagged as potential doublets and excluded from further analysis. Quality control thresholds were then applied to retain high-quality cells: we required each cell to have >300 detected genes and a mitochondrial gene content <30% (to exclude dying cells with high mitochondrial RNA). Cells failing these criteria were removed. After QC filtering, gene expression counts were normalized to counts per 10,000, log-transformed, and highly variable genes were identified for each sample.

### Data integration and clustering

To integrate single-cell data across the 15 samples while accounting for batch effects, we performed joint analysis via batch correction and graph-based clustering. The per-sample Scanpy AnnData objects were concatenated, and we used batch-balanced k-nearest neighbors (BBKNN)^94^ to correct for batch effects across different patients. Specifically, a PCA (principal component analysis) was run on the combined data (using the top 50 principal components), and BBKNN was applied with sample_batch as the batch key to align the data in the PCA space. This method retains biological variability while mitigating technical batch differences. We then computed a two-dimensional uniform manifold approximation and projection (UMAP) embedding for visualization of the integrated dataset. Clustering of cells was performed with the Leiden graph-based community detection algorithm^95^ (Scanpy) via the 50 PCs (principal components) and the BBKNN-adjusted neighbor graph. As a result, the cells were partitioned into distinct clusters corresponding to putative cell types or states. Each cluster was examined in the context of known cell type markers for annotation.

### Cell type annotation

We assigned cell identities to clusters via two complementary automated tools. CellO (cell ontology-based classifier)^11^ was applied to predict cell types hierarchically on the basis of Cell Ontology. CellO uses a machine learning model trained on a broad compendium of human cell types and provides probabilistic, ontology-informed annotations for each cell. In parallel, we used CellTypist for rapid cell labeling. CellTypist^12^ uses logistic regression classifiers (optimized via stochastic gradient descent) and comes with built-in models (focused especially on immune cell subtypes) for automated cell type prediction. The combination of CellO and CellTypist helped refine the annotations: CellO provided a hierarchical view of cell identity, whereas CellTypist offered high-speed predictions with curated models. The final cell type labels for each cluster were determined by cross-referencing the automated predictions with known marker gene expression in our data.

### Copy number variation inference

To distinguish malignant (aneuploid) epithelial cells from normal diploid cells, we inferred large-scale copy number variations from the transcriptome data via CopyKAT (copy number karyotyping of tumors)^14^. CopyKAT applies an integrative Bayesian approach to estimate genome-wide copy number profiles at ∼5 Mb resolution from scRNA-seq gene expression data, enabling the identification of aneuploid subpopulations. The underlying rationale is that groups of adjacent genes show concerted expression changes when their genomic region is amplified or deleted; by analyzing these patterns, CopyKAT segregates tumor cells (with widespread aneuploidy) from stromal or immune cells, which typically have near-diploid profiles. We ran CopyKAT in R (v1.1.0) on log-normalized expression data for epithelial compartments, which yielded a copy number heatmap and classified each cell as “aneuploid-tumor” or “diploid” along with the identification of possible tumor subclones.

### Neutrophil differential gene expression and regulatory analysis

To obtain replicate-aware gene-expression profiles, raw counts from all neutrophils belonging to the same biological replicate (patient) were combined into pseudobulk profiles using decoupler v1.4.0^54^. Replicates were stratified into two clinical groups: control/non-invasive (benign endometrium + EIN) and invasive (transition + EC). Pseudobulk count matrices were analyzed with DESeq2 v1.36.0^96^. Library sizes were internally normalized with the median-of-ratios method; dispersions were estimated per gene; and model fitting used the Wald test, specifying group as the single design factor. The log₂ fold-changes were shrunk with the apeglm estimator to stabilize the variance for genes with low expression. Genes with Benjamini–Hochberg–adjusted *p* < 0.05 and |log₂FC| > 0.5 were deemed significantly differentially expressed (DEGs). Effect sizes and adjusted *p*-values were visualized in volcano plots; the twenty most significant DEGs were annotated for reference.

To infer upstream regulation, we applied the unified linear model (ULM) framework implemented in the decoupler. Gene-level Wald statistics from the DESeq2 contrast (invasive vs. control) were regressed against the CollecTRI regulon compendium^67^, (human, non-split complexes; >16,000 each transcription factor (TF)–target interactions collated from 12 databases). The ULM returns an activity score and empirical *p*-value for TFs; TFs with a false-discovery-rate (FDR) < 0.05 were considered significantly activated or repressed in invasive neutrophils.

The functional interpretation of the DEGs was performed via over-representation analysis (ORA) against the MSigDB hallmark gene-set collection (v2023.2)^52^. Significant DEGs (*padj* < 0.05) were tested for enrichment via one-tailed Fisher’s exact tests, with FDR correction across 50 hallmark terms. Enriched pathways (FDR < 0.05) are summarized in dot plots showing the combined enrichment score, odds ratio, and pathway FDR.

### Stage-correlated GO terms

To pinpoint biological processes whose transcriptional activity tracks with histologic progression from a benign endometrium to carcinoma, we applied a MAST^16^-based gene-set regression strategy to single-cell profiles. Each cell carried a four-level ordered factor denoting the disease stage—control, EIN, transition, or EC—which was additionally converted to a numeric covariate (1 to 4) for monotonic-trend testing. For each gene, MAST’s two-part-hurdle model, log-link for positive counts and logistic for detection, was fitted with the numeric stage variable as the sole fixed effect and with the combined-residuals hook to adjust for the cellular detection rate. Model coefficients and their variance–covariance matrices were stabilized by non-parametric bootstrap resampling, providing gene-wise Wald Z statistics for the stage term. GO Biological Process (2023)^29,30^ gene sets were downloaded from EnrichR^97^, mapped to expressed genes, and filtered to modules containing ≥ 5 genes. Using gseaAfterBoot, per-gene stage *Z* scores were aggregated into set-level statistics that incorporate bootstrap-derived covariance, yielding a combined *Z* and adjusted *p*-value (Benjamini–Hochberg) for each module. Sets with FDR < 1 × 10⁻⁴ were deemed significantly correlated (positively or negatively) with EC progression; the top 50 modules by |combined Z| were retained for visualization. For representative GO modules, single-cell module scores were generated with Seurat’s^98^ AddModuleScore (100 control genes per module). Stage-wise distributions are displayed as violin plots overlaid with LOESS-smoothed mean trajectories.

### RNA velocity

We leveraged scVelo^13^ to calculate the RNA velocity vectors for each cell. scVelo generalizes the RNA velocity method by modeling splicing kinetics, enabling the estimation of future transcriptional states of cells without assuming a steady state. Briefly, we obtained spliced and unspliced transcript counts from the Cell Ranger^92^ output and then ran scVelo (v0.2.4) in dynamical mode to determine gene-specific transcription and splicing rates. This yielded a “velocity field” on the UMAP, indicating the direction of state transitions for cells. From the scVelo model, we also extracted a latent time ordering of cells, which is a data-driven pseudotemporal timeline consistent with RNA velocity. Together, these analyses allowed us to characterize the continuum of cellular state changes during EC progression and to identify potential driver genes whose up- or downregulation drives the transition via scVelo’s likelihood-based ranking of putative dynamic genes.

### Gene-set enrichment analysis

We applied gene set enrichment analysis (GSEA)^76^ for ranked gene lists ordered by log-fold change between two stages with gseapy^99^ (Python implementation of GSEApy v1.1.0). Normalized enrichment scores (NESs), nominal *P* values, and FDR-adjusted *q* values (Benjamini– Hochberg) are reported. Pathways with |NES| > 1.5 and FDR < 0.05 were considered significantly positively or negatively enriched. For discrete DEG lists, over-representation analysis (ORA) was carried out with gprofiler2^100^. The query genes were mapped to Ensembl identifiers and tested against the g:GOSt catalog, which includes Gene Ontology (BP, MF, CC) ^29,30^, KEGG^86^, and Reactome^80^ pathways. The built-in g:SCS multiple-testing correction was applied; terms with FDR < 0.05 were retained. The enrichment outputs—odds ratios, adjusted *p*-values, and term sizes—were integrated into a dot-plot.

### Temporal expression dynamics

We utilized TDEseq^82^ (temporal differential expression sequencing) to rigorously model gene expression as a function of EC development. TDEseq is a nonparametric method designed for multisample, multistage scRNA-seq data that uses smoothing splines to account for time/stage dependencies and mixed-effect models to handle intrasample correlations. This approach classifies genes into distinct temporal patterns (e.g., monotonically increasing, decreasing) across the ordered stages. We applied TDEseq (R package v1.1) to neutrophils and used the four stages as ordered factors. The analysis yielded lists of stage-dynamic genes and their assigned temporal pattern categories.

### State classification modeling

We trained a stochastic gradient descent classifier (implemented via scikit-learn’s^101^ SGDClassifier) to classify cells into their state of origin (cancer-free, EIN, transition, early EC) on the basis of their normalized gene expression features. The classifier was configured as a logistic regression (with elastic net regularization) optimized by SGD. Model performance was assessed via cross-validation on held-out cells (33% of the cells that were not used in the training process) and summarized with confusion matrices (true state vs. predicted state). Important features (genes) contributing to the classification were extracted from the model coefficients.

## Data and code availability

Data and code available upon request.

## Acknowledgment

We thank Laura Hansen, Murray Casey, and James Grunkemeyer for their valuable input in the establishment of this project. We thank the Creighton University Innovative Genomics Core for the assistance.

## Author contributions

Conceptualization: L.C., R.F., Y.F.

Methodology: L.C., S.Y., L.K., H.B., K.A., J.C., R.F., J.X., Y.F.

Data analysis and visualization: B.F., Y.F.

Project administration: L.C., J.X., Y.F.

Writing: L.C., J.C., J.X., Y.F.

## Supplementary materials

Figures S1 to S9

Table S1. Demographic and clinical characteristics of participants.

