## Supplementary material for "Dynamic uterine microenvironment drives endometrial adenocarcinoma carcinogenesis and progression": Figures S1 to S9

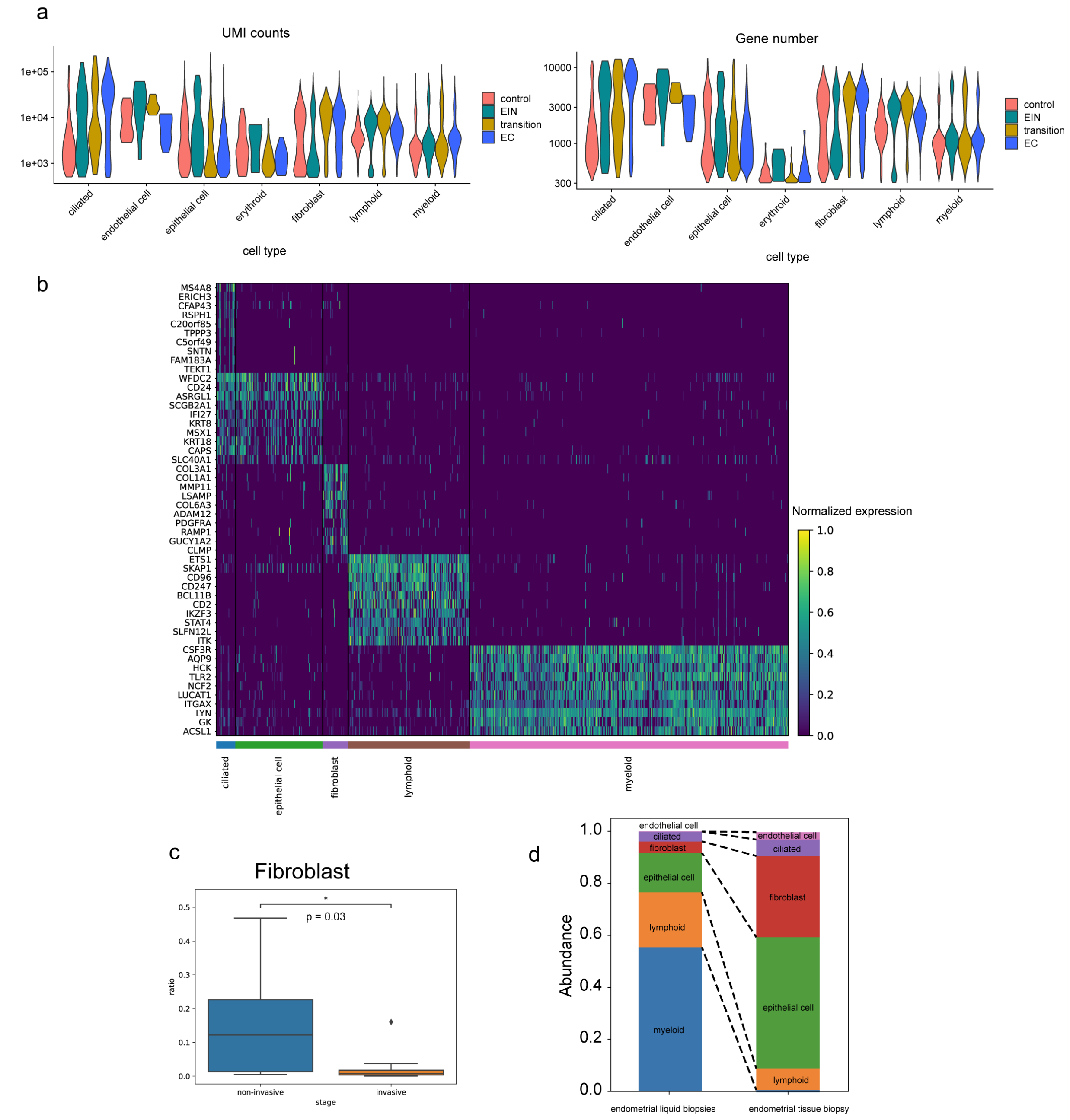


**Supplemental Figure S1. Quality control and abundance changes across EC progression**

(a) Violin plots of UMI counts (left) and gene numbers (right) for each major cell type across the 4 disease stages.

(b) Heatmap of cell type-specific marker gene expression across all major cell types.

(c) Box plot showing differences in the fibroblast ratio between noninvasive and invasive ECs. Statistical tests revealed a significant reduction in fibroblasts in invasive cases (Mann–Whitney $p = 0.03$).

(d) Stacked bar plot comparing the relative abundance of major cell types in matched endometrial liquid biopsies (left) and endometrial tissue biopsies (right). The bars represent the proportional composition of six cell classes (myeloid, lymphoid, epithelial, fibroblast, ciliated epithelial, and endothelial), with dashed lines connecting equivalent cell types across biopsy types.

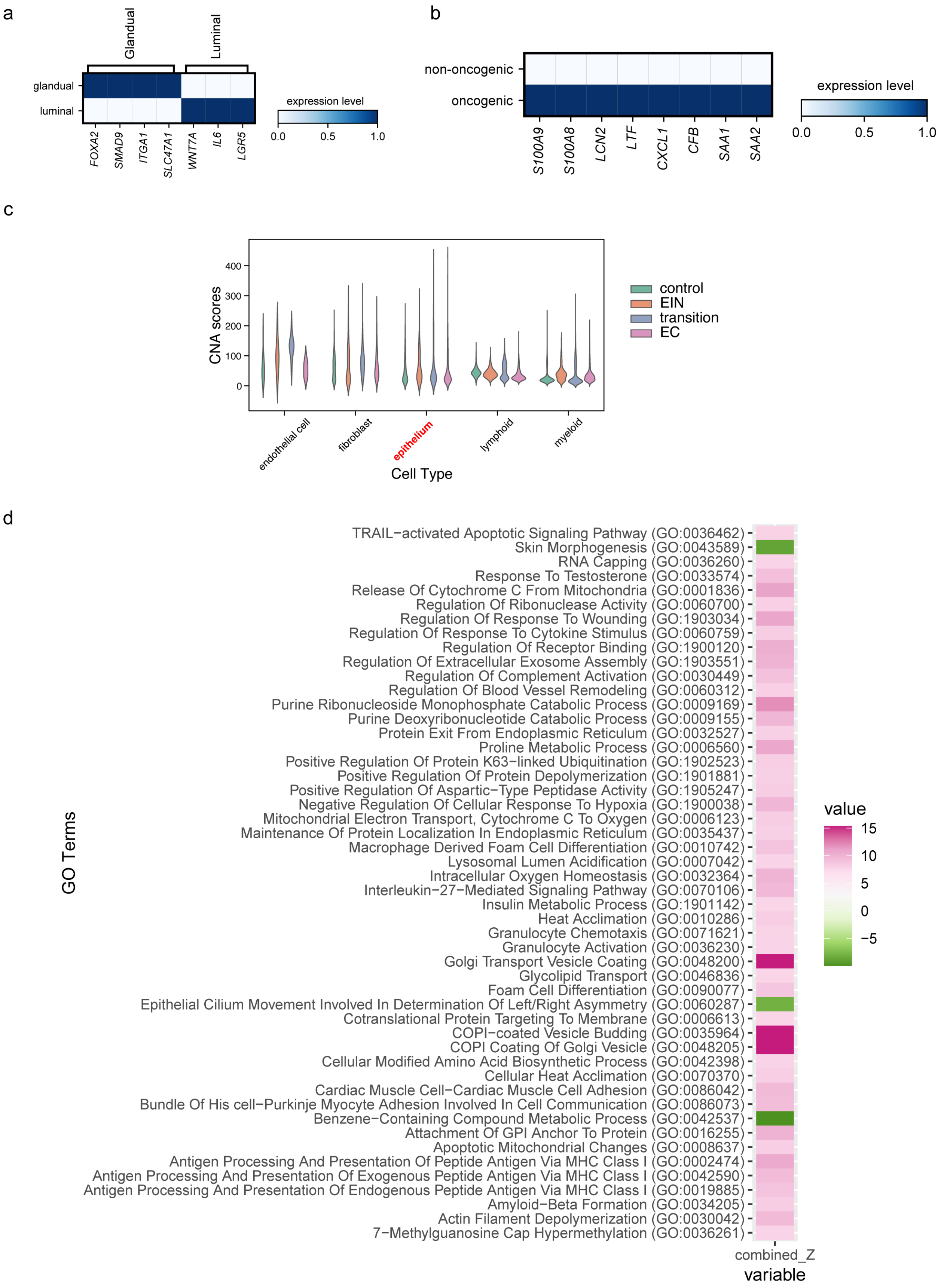

a
Glandual
Luminal
glandual
luminal
FOXA2
SMAD9
ITGA1
SLC47A1
WNT7A
IL6
LGR5
expression level
0.0
0.5
1.0
b
non-oncogenic
oncogenic
S100A9
S100A8
LCN2
LTF
CXCL1
CFB
SAA1
SAA2
expression level
0.0
0.5
1.0
c
CNA scores
0
100
200
300
400
endothelial cell
fibroblast
epithelium
lymphoid
myeloid
Cell Type
control
EIN
transition
EC
d
GO Terms
TRAIL−activated Apoptotic Signaling Pathway (GO:0036462)
Skin Morphogenesis (GO:0043589)
RNA Capping (GO:0036260)
Response To Testosterone (GO:0033574)
Release Of Cytochrome C From Mitochondria (GO:0001836)
Regulation Of Ribonuclease Activity (GO:0060700)
Regulation Of Response To Wounding (GO:1903034)
Regulation Of Response To Cytokine Stimulus (GO:0060759)
Regulation Of Receptor Binding (GO:1900120)
Regulation Of Extracellular Exosome Assembly (GO:1903551)
Regulation Of Complement Activation (GO:0030449)
Regulation Of Blood Vessel Remodeling (GO:0060312)
Purine Ribonucleoside Monophosphate Catabolic Process (GO:0009169)
Purine Deoxyribonucleotide Catabolic Process (GO:0009155)
Protein Exit From Endoplasmic Reticulum (GO:0032527)
Proline Metabolic Process (GO:0006560)
Positive Regulation Of Protein K63−linked Ubiquitination (GO:1902523)
Positive Regulation Of Protein Depolymerization (GO:1901881)
Positive Regulation Of Aspartic−Type Peptidase Activity (GO:1905247)
Negative Regulation Of Cellular Response To Hypoxia (GO:1900038)
Mitochondrial Electron Transport, Cytochrome C To Oxygen (GO:0006123)
Maintenance Of Protein Localization In Endoplasmic Reticulum (GO:0035437)
Macrophage Derived Foam Cell Differentiation (GO:0010742)
Lysosomal Lumen Acidification (GO:0007042)
Intracellular Oxygen Homeostasis (GO:0032364)
Interleukin−27−Mediated Signaling Pathway (GO:0070106)
Insulin Metabolic Process (GO:1901142)
Heat Acclimation (GO:0010286)
Granulocyte Chemotaxis (GO:0071621)
Granulocyte Activation (GO:0036230)
Golgi Transport Vesicle Coating (GO:0048200)
Glycolipid Transport (GO:0046836)
Foam Cell Differentiation (GO:0090077)
Epithelial Cilium Movement Involved In Determination Of Left/Right Asymmetry (GO:0060287)
Cotranslational Protein Targeting To Membrane (GO:0006613)
COPI−coated Vesicle Budding (GO:0035964)
COPI Coating Of Golgi Vesicle (GO:0048205)
Cellular Modified Amino Acid Biosynthetic Process (GO:0042398)
Cellular Heat Acclimation (GO:0070370)
Cardiac Muscle Cell−Cardiac Muscle Cell Adhesion (GO:0086042)
Bundle Of His cell−Purkinje Myocyte Adhesion Involved In Cell Communication (GO:0086073)
Benzene−Containing Compound Metabolic Process (GO:0042537)
Attachment Of GPI Anchor To Protein (GO:0016255)
Apoptotic Mitochondrial Changes (GO:0008637)
Antigen Processing And Presentation Of Peptide Antigen Via MHC Class I (GO:0002474)
Antigen Processing And Presentation Of Exogenous Peptide Antigen Via MHC Class I (GO:0042590)
Antigen Processing And Presentation Of Endogenous Peptide Antigen Via MHC Class I (GO:0019885)
Amyloid−Beta Formation (GO:0034205)
Actin Filament Depolymerization (GO:0030042)
7−Methylguanosine Cap Hypermethylation (GO:0036261)
combined_Z
variable
value
15
10
5
0
−5

**Supplemental Figure S2. Validation of epithelial cell subpopulation and gene module enrichment across EC development**

(a) Heatmap of glandular and luminal epithelial marker gene expression confirming accurate annotation of unciliated epithelial subtypes in uterine blood-derived cells.

(b) Expression levels of oncogenic markers are enriched in a distinct epithelial subpopulation.

(c) Violin plots of copy number alteration (CNA) scores across major cell types and EC stages reveal a sharp increase in CNA burden specific to epithelial cells during progression.

(d) Heatmap of the top 50 Gene Ontology (GO) terms correlated with EC development stages, with scores reflecting their relationship with disease progression.

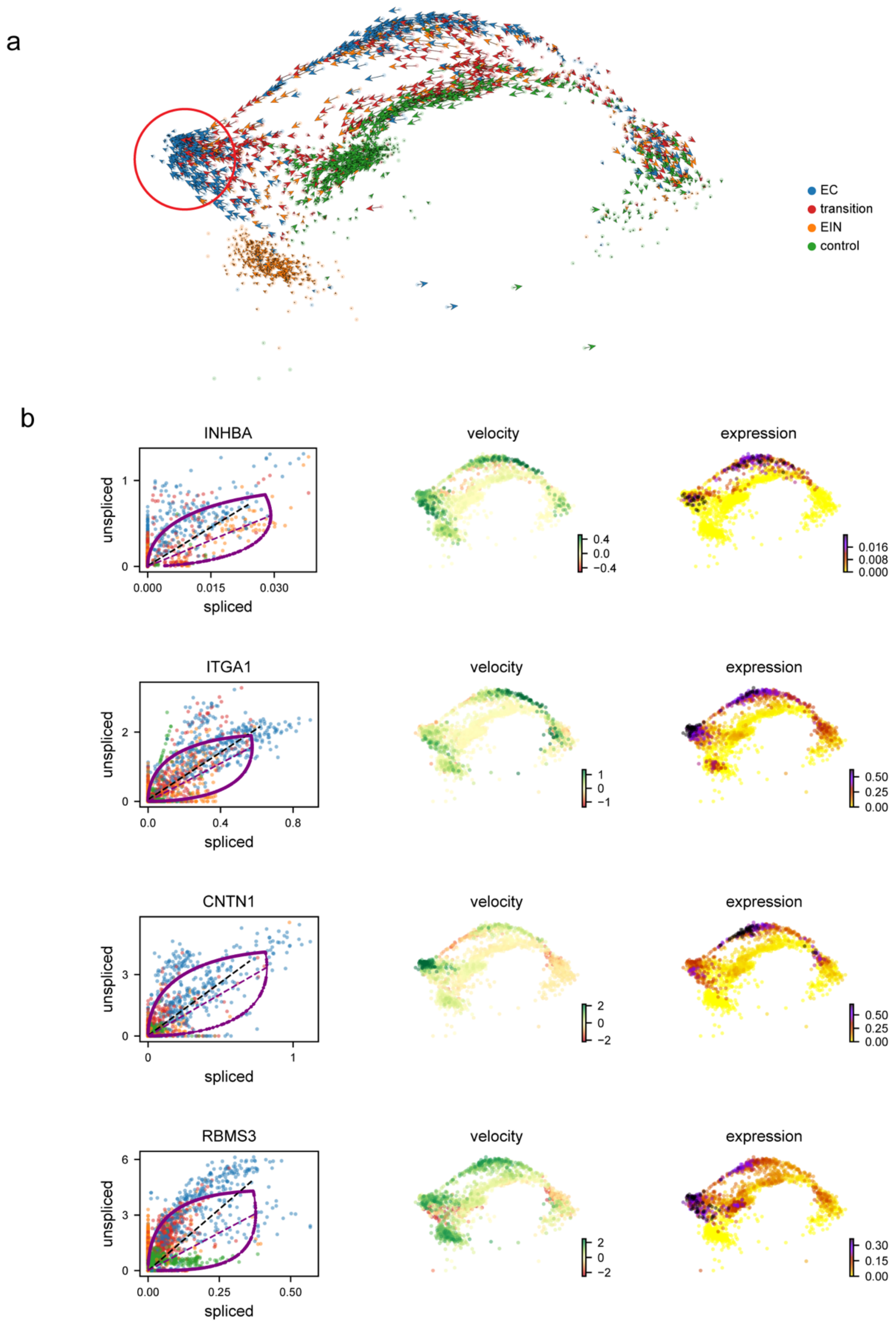


**Supplemental Figure S3. RNA-velocity dissects the step-wise re-programming of fibroblasts during EC progression**

(a) UMAP of all fibroblasts according to pathological stage—green, control; orange, EIN; red, transition; blue, EC. Arrows represent RNA-velocity vectors predicted with the dynamical model in scVelo; arrow length encodes the speed and direction of the inferred future transcriptional state. Vectors converge from a control-enriched root toward an EC-dominant terminus (red circle), indicating a continuous developmental trajectory from normal to cancer-associated fibroblasts.

(b) Gene-specific velocity phase portraits and embedding overlays for the top velocity drivers. For each gene, three panels are shown: phase portrait (left): unsliced (y-axis) versus spliced (x-axis) counts for individual cells, colored by stage as in *a*. The fitted latent-time trajectory (purple curve) denotes transcriptional dynamics; dashed arrows indicate the velocity direction in phase space. Velocity embedding (middle): UMAP of fibroblasts colored by cell-wise velocity values for the indicated gene (green = positive induction, brown = negative). Expression embedding (right): log-normalized expression of the same gene projected onto UMAP (yellow/purple gradient; scale bars show the z-scores).

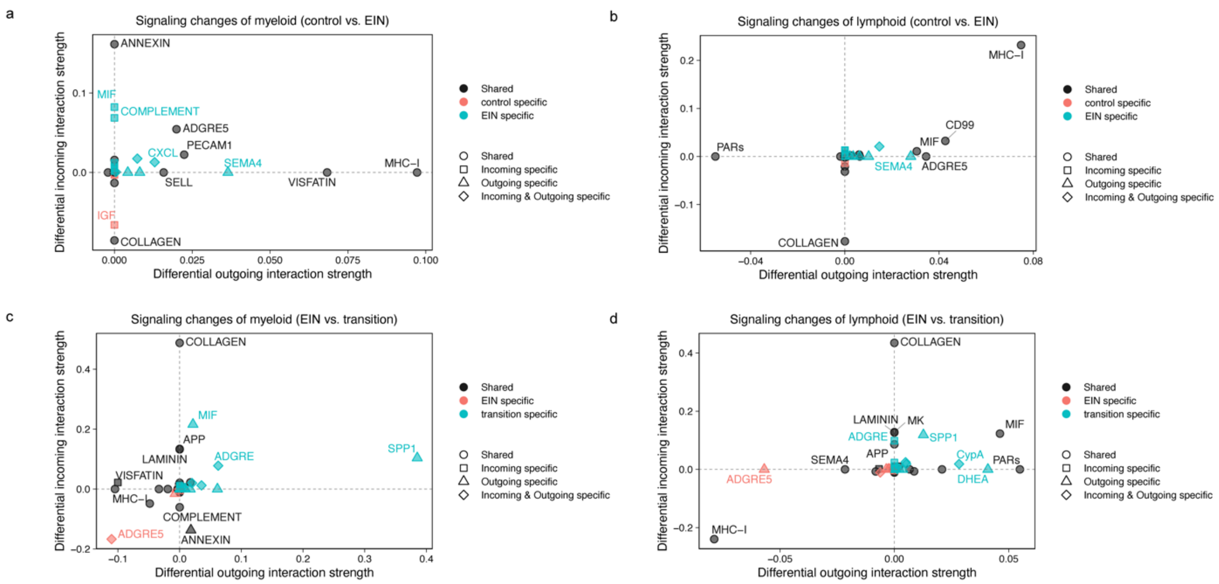


**Supplemental Figure S4. Differential ligand–receptor signaling changes in myeloid and lymphoid cells during EC development**

(a–b) Scatter plots showing differential incoming and outgoing interaction strengths of signaling pathways in myeloid (a) and lymphoid (b) cells between the control and EIN groups.

(c–d) Similar analyses comparing EIN vs. transition for myeloid (c) and lymphoid (d) cells. Dot shape represents interaction specificity (incoming, outgoing, or both); dot color indicates group specificity.

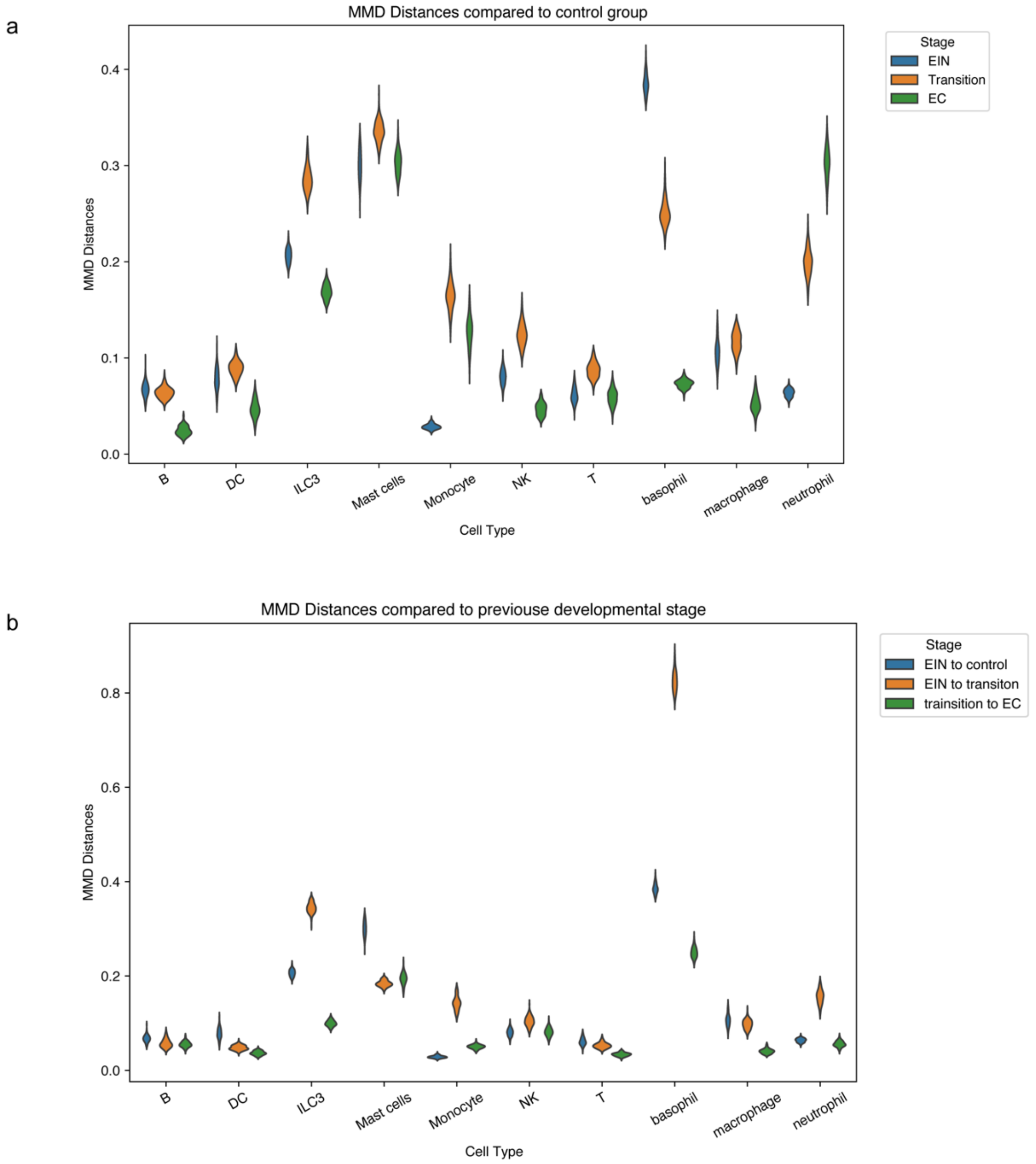


**Supplemental Figure S5. MMD analysis of immune-cell state changes**

(a) Violin plots of MMD distances between each immune cell type at the indicated stages and its control counterpart (larger values = greater divergence).

(b) Violin plots of MMD distances between consecutive stages (EIN→control, EIN→transition, transition→EC), representing the instantaneous “speed” of transcriptional change.

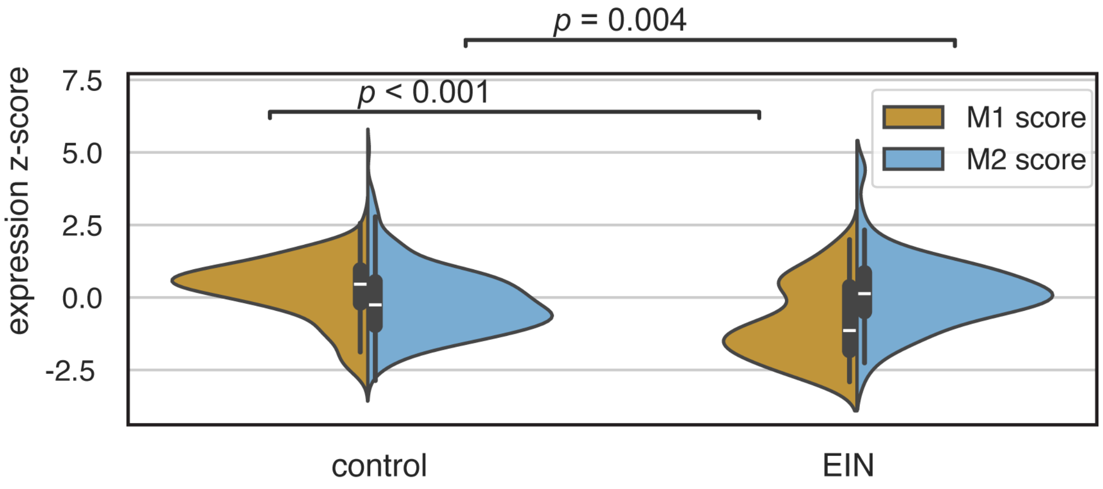


**Supplemental Figure S6. Progressive immune reprogramming during EC development.** Violin plots of classical- (M1, ochre) and alternative- (M2, blue) macrophage signature scores in control versus EIN samples. Centre lines mark medians; whiskers denote 1.5× IQR. *P* values were calculated from two-sided Wilcoxon tests.

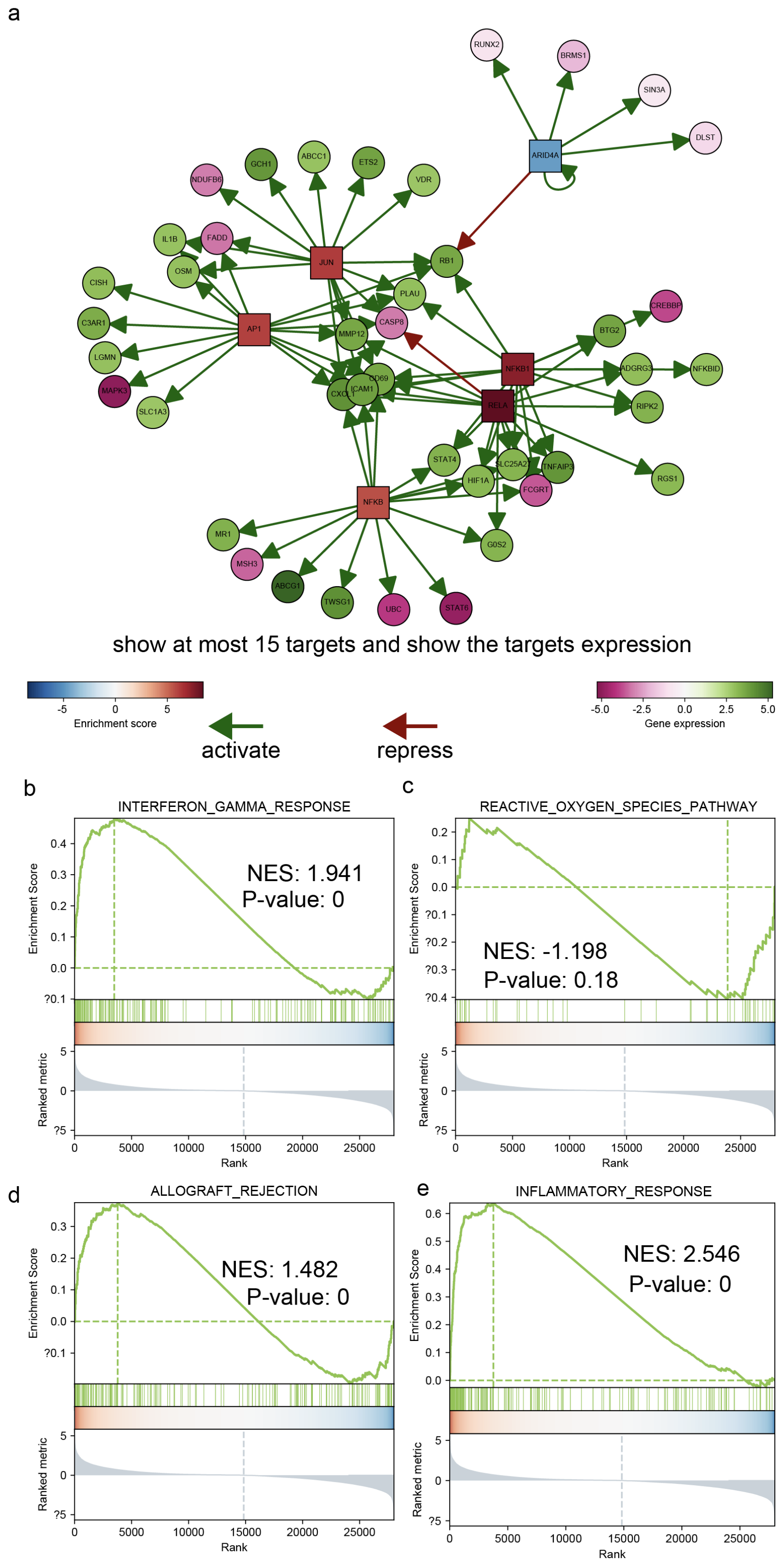

a
RUNX2
BRMS1
SIN3A
DLST
ARID4A
NDUFB6
GCH1
ABCC1
ETS2
VDR
IL1B
FADD
OSM
JUN
RB1
CISH
PLAU
C3AR1
AP1
CASP8
CREBBP
LGMN
MMP12
BTG2
MAPK3
CD69
NFKB1
ADGRG3
NFKBID
CXCL
ICAM1
SLC1A3
RELA
RIPK2
STAT4
SLC25A2
TNFAIP3
RGS1
HIF1A
FCGRT
NFKB
MR1
G0S2
MSH3
ABCG1
TWSG1
UBC
STAT6
show at most 15 targets and show the targets expression
-5
0
5
Enrichment score
-5.0
-2.5
0.0
2.5
5.0
Gene expression
activate
repress
b
INTERFERON_GAMMA_RESPONSE
NES: 1.941
P-value: 0
Enrichment Score
Ranked metric
Rank
c
REACTIVE_OXYGEN_SPECIES_PATHWAY
NES: -1.198
P-value: 0.18
Enrichment Score
Ranked metric
Rank
d
ALLOGRAFT_REJECTION
NES: 1.482
P-value: 0
Enrichment Score
Ranked metric
Rank
e
INFLAMMATORY_RESPONSE
NES: 2.546
P-value: 0
Enrichment Score
Ranked metric
Rank

**Supplemental Figure S7. Regulatory and pathway landscape of invasive neutrophils**

(a) CollecTRI interaction graph illustrating activated (green arrows) or repressed (red arrows) targets of key TFs; node shading represents $log_2FC$. TF shading represents the enrichment score.

(b–e), GSEA enrichment curves for the interferon-γ response (b), the ROS pathway (c), allograft rejection (d) and the inflammatory response (e); NES and *P*-values are indicated.

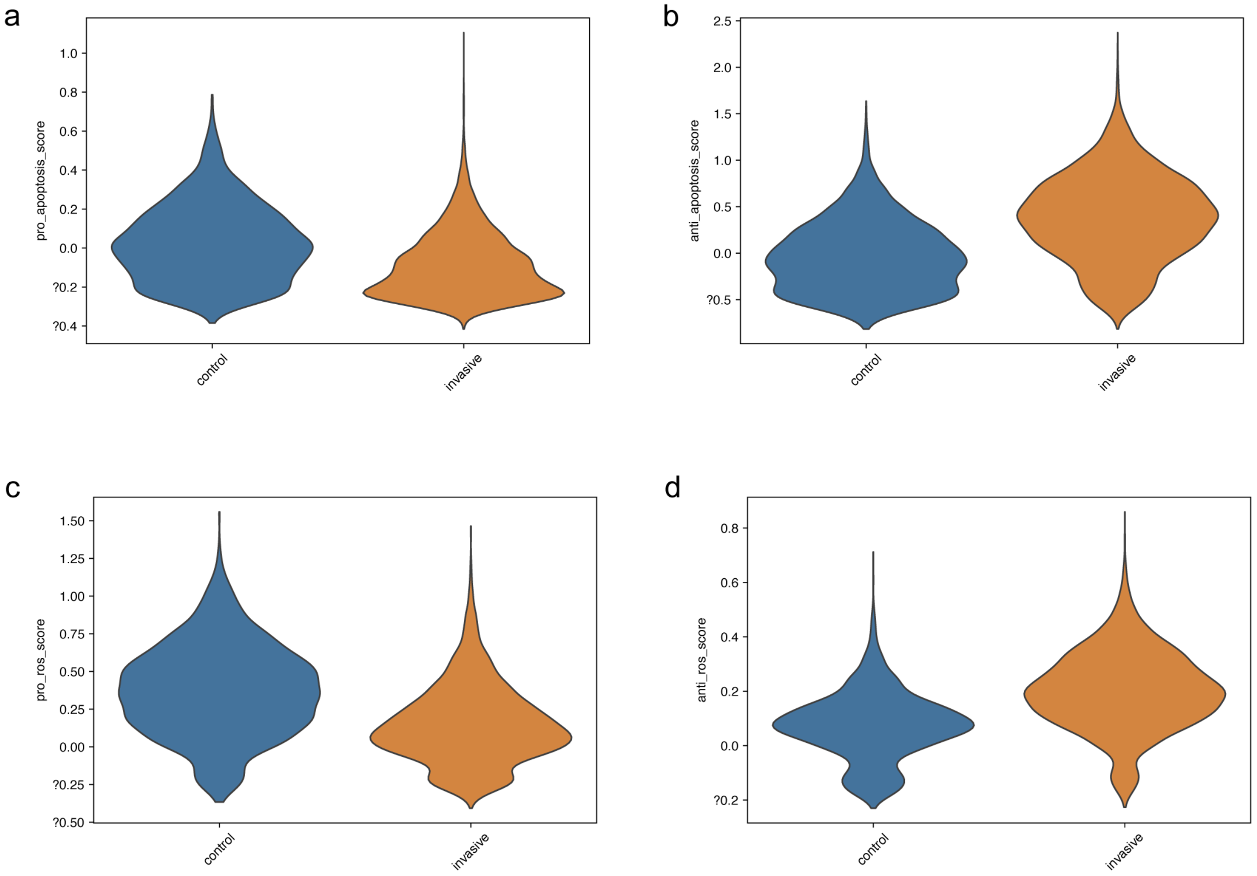


**Supplemental Figure S8. Distributions of ROS- and apoptosis-related module scores**

**(a–d)** Violin plots comparing pro- and anti-apoptotic (a, b) and pro- and anti-ROS (c, d) signatures between non-invasive and invasive neutrophils.

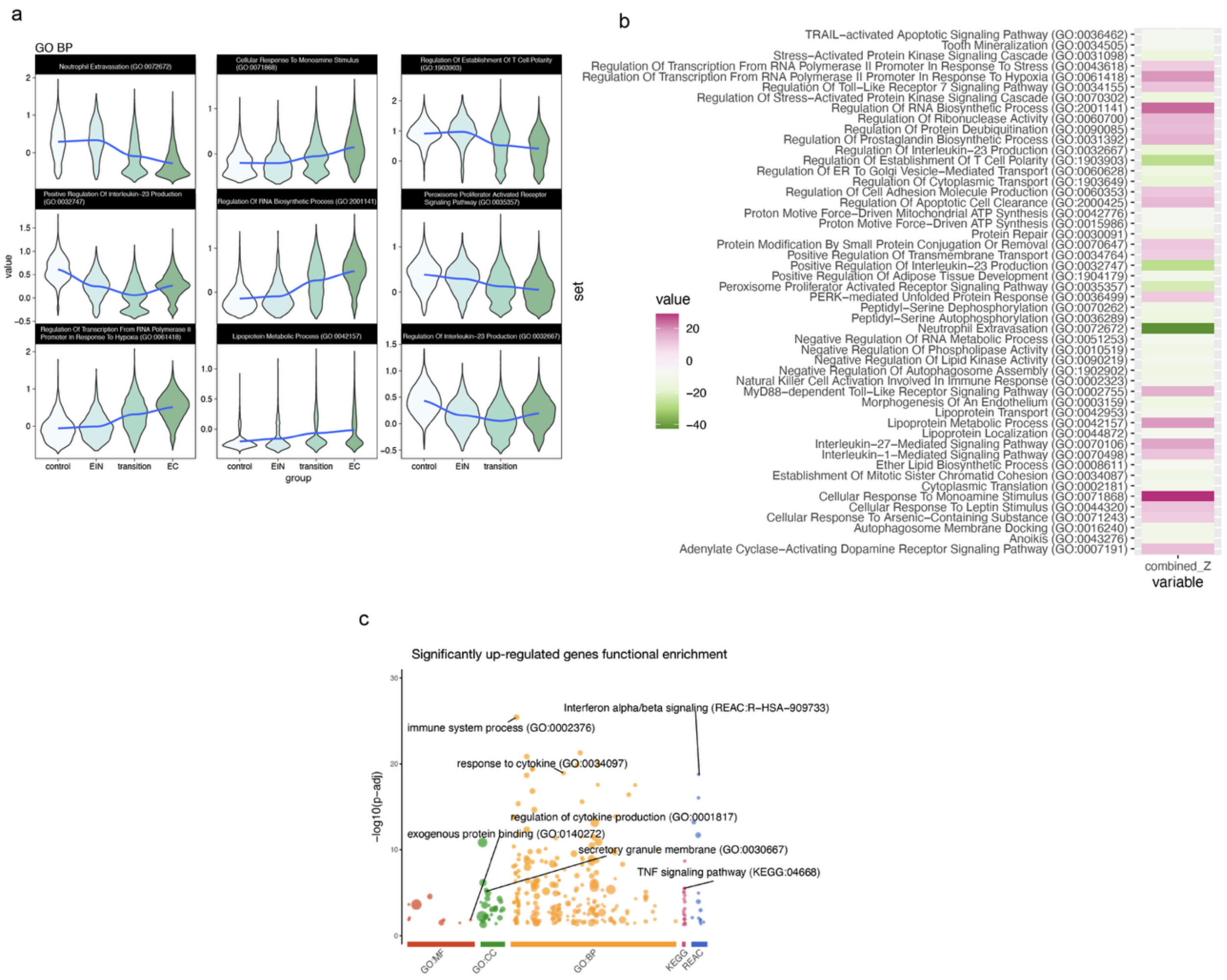

**Supplemental Figure S9. Functional annotation of stage-correlated neutrophils**

(a–b) Violin plots (a) and heat-map (b) highlighting GO terms whose module scores correlate positively or negatively with pathological stage.

(c) Bubble plot of enriched GO and pathway terms for EIN-up-regulated genes; bubble size = gene count, color = $-\log_{10}$ adjusted P.
